# SWI/SNF senses carbon starvation with a pH-sensitive low complexity sequence

**DOI:** 10.1101/2021.03.03.433592

**Authors:** J. Ignacio Gutiérrez, Gregory P. Brittingham, Yonca B. Karadeniz, Kathleen D. Tran, Arnob Dutta, Alex S. Holehouse, Craig L. Peterson, Liam J. Holt

## Abstract

It is increasingly appreciated that intracellular pH changes are important biological signals. This motivates the elucidation of molecular mechanisms of pH-sensing. We determined that a nucleocytoplasmic pH oscillation was required for the transcriptional response to carbon starvation in *S. cerevisiae*. The SWI/SNF chromatin remodeling complex is a key mediator of this transcriptional response. We found that a glutamine-rich low complexity sequence (QLC) in the *SNF5* subunit of this complex, and histidines within this sequence, were required for efficient transcriptional reprogramming during carbon starvation. Furthermore, the *SNF5* QLC mediated pH-dependent recruitment of SWI/SNF to a model promoter *in vitro.* Simulations showed that protonation of histidines within the *SNF5* QLC lead to conformational expansion, providing a potential biophysical mechanism for regulation of these interactions. Together, our results indicate that that pH changes are a second messenger for transcriptional reprogramming during carbon starvation, and that the *SNF5* QLC acts as a pH-sensor.

## Introduction

Biological processes are inherently sensitive to the solution environment in which they occur. A key regulated parameter is intracellular pH (pH_i_), which influences all biological processes by determining the protonation state of titratable chemical groups. These titratable groups are found across many biological molecules, from small-molecule osmolytes to the side-chains of amino acids. While early work suggested that pH_i_ was a tightly constrained cellular parameter (1), more recent technologies have revealed that pH_i_ can vary substantially in both space and time (2, 3). Moreover changes in pH_i_ can regulate metabolism (4, 5), proliferation (6), and cell fate (7), among other processes. Intriguingly, stress-associated intracellular acidification appears to be broadly conserved, suggesting that a drop in pH_i_ is a primordial mechanism to coordinate the general cellular stress response (8–13).

The budding yeast *Saccharomyces cerevisiae* is adapted to an acidic external environment (pH_e_), and optimal growth media is typically at pH 4.0 – 5.5. The plasma membrane (Pma1) and vacuolar (Vma1) ATPases maintain near neutral pH_i_ of ∼7.8 by pumping protons out of the cell and into the vacuole, respectively (14). When cells are starved for carbon, these pumps are inactivated, leading to a rapid acidification of the intracellular space to pH ∼ 6 (15, 16). This decrease in intracellular pH_i_ is crucial for viability upon carbon-starvation, and is thought to conserve energy, leading to storage of metabolic enzymes in filamentous assemblies (17), reduction of macromolecular diffusion (18, 19), decreased membrane biogenesis (4) and possibly the non-covalent crosslinking of the cytoplasm into a solid-like material state (18, 19). These studies suggest that many physiological processes are inactivated when pH_i_ drops. However, some processes must also be upregulated during carbon starvation to enable adaptation to this stress. These genes are referred to as “glucose-repressed genes”, as they are transcriptionally repressed in the presence of glucose (20, 21). Recently, evidence was presented of a positive role for acidic pH_i_ in stress-gene induction: transient acidification is required for induction of the transcriptional heat-shock response in some conditions (13). However, the molecular mechanisms by which the transcriptional machinery senses and responds to pH changes remain mysterious.

The <u>S</u>ucrose <u>N</u>on <u>F</u>ermenting genes (*SNF*) were among the first genes found to be required for induction of glucose-repressed genes (22). Several of these genes were later identified as members of the SWI/SNF complex (23, 24), an 11 subunit chromatin remodeling complex that is highly conserved from yeast to mammals (25–27). The SWI/SNF complex affects the expression of ∼10% of the genes in *Saccharomyces cerevisiae* during vegetative growth (28). Upon carbon starvation, most genes are down-regulated, but a set of glucose-repressed genes, required for utilization of alternative energy sources, are strongly induced (21). The SWI/SNF complex is required for the efficient expression of several hundred stress-response and glucose-repressed genes, implying a possible function in pH-associated gene expression (28, 29). However, we still lack evidence for a direct role for SWI/SNF components in the coordination of pH-dependent transcriptional programs or a mechanism through which pH-sensing may be achieved.

10/11 subunits of the SWI/SNF complex contain large intrinsically disordered regions (**Figure 1 – figure supplement 1**), and in particular, 4/11 SWI/SNF subunits contain glutamine-rich low complexity (QLC) sequences. QLCs are present in glutamine-rich transactivation domains (30, 31) some of which, including those found within SWI/SNF, may bind to transcription factors (32), or recruit transcriptional machinery (33–35). Intrinsically disordered regions lack a fixed three dimensional structure and have been proposed to be highly responsive to their solution environment (36, 37). Moreover, the SWI/SNF QLCs contain multiple histidine residues. Given that the intrinsic pK_a_ of the histidine sidechain is 6.9 (38), we hypothesized that these glutamine-rich low complexity regions might function as pH sensors in response to variations in pH_i._

In this study, we elucidate *SNF5* as a pH-sensing regulatory subunit of SWI/SNF. *SNF5* is over 50% disordered and contains the largest QLC of the SWI/SNF complex. This region is 42% glutamine and contains 7 histidine residues. We investigated the relationship between the *SNF5* QLC and the cytosolic acidification that occurs during acute carbon-starvation. By single cell analysis, we found that intracellular pH (pH_i_) is highly dynamic and varies between subpopulations of cells within the same culture. After an initial decrease to pH_i_ ∼ 6.5, a subset of cells recovered their pH_i_ to ∼ 7. This transient acidification followed by recovery was required for expression of glucose-repressed genes. The *SNF5* QLC and four embedded histidines were required for rapid gene induction. SWI/SNF complex histone remodeling activity was robust to pH changes, but recruitment of the complex to a model transcription factor was pH-sensitive, and this recruitment was mediated by the *SNF5* QLC. All-atom simulations indicated that histidine protonation causes a conformational expansion of the *SNF5* QLC, perhaps enabling interaction with a different set of transcription factors and driving recruitment to the promoters of glucose-repressed genes. Thus, we propose changes in histidine charge within QLCs as a mechanism to sense pH changes and instruct transcriptional reprograming during carbon starvation.

## Results

### Induction of *ADH2* upon glucose starvation requires the *SNF5* glutamine-rich low complexity sequence with native histidines

The SWI/SNF chromatin remodeling complex subunit *SNF5* has a large low-complexity region at its N-terminus that is enriched for glutamine, the sequence of which is shown in figure 1A. This sequence contains seven histidine residues, and we noticed a frequent co-occurrence of histidines within and adjacent to glutamine-rich low complexity sequences (QLCs) of many proteins. Inspection of the sequence properties of proteins, especially through the lens of evolution, can provide hints as to functionally important features. Therefore, we analyzed the sequence properties of all glutamine-rich low complexity sequences (QLCs) in the proteomes of several species. We defined QLCs as stretches of low-complexity sequence containing at least 10 glutamines. We allowed interruption of the glutamines with any number of single or double amino acid insertions, but a QLC was terminated by an interruption of 3 or more non-Q amino acids (see methods). By these criteria, the S288c *S. cerevisiae* strain had 116 QLCs (**Supplemental Table 1**). We found that Alanine, Proline and Histidine were enriched (> 2-fold higher than average proteome abundance) in yeast QLCs (Figure 1B), with similar patterns found in *Dictyostelium discoidum*, and *Drosophila melanogaster* proteomes (**Figure 1 – figure supplement 2**). Enrichment for histidine within QLCs was previously described across many *Eukaryotes* using a slightly different method (39). Interestingly, the codons for glutamine are a single base pair mutation away from proline and histidine. However, they are similarly adjacent to lysine, arginine, glutamate and leucine, yet QLCs are depleted for lysine, arginine and glutamate, suggesting that the structure of the genetic code is insufficient to explain the observed patterns of amino acids within QLCs. We also considered the possibility that histidines might be generally enriched in low-complexity sequences. In fact, this is not the case: histidines are 7-fold more abundant in yeast QLCs than in all low-complexity sequences identified using Wooton-Fedherhen complexity (see methods). Thus, histidines are a salient feature of QLCs.

**Figure 1:**
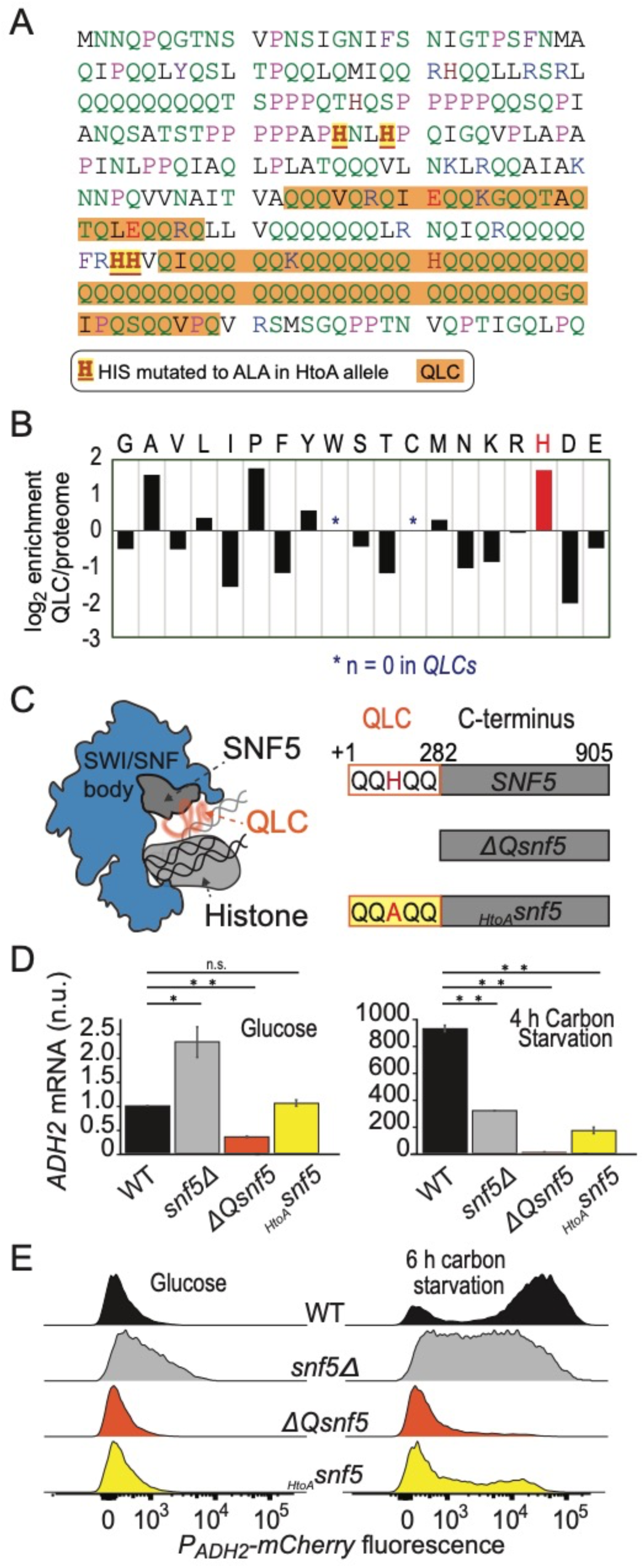
Efficient induction of *ADH2* upon glucose starvation requires the *SNF5* glutamine-rich low complexity sequence with native histidines. **A)** Sequence of the N-terminal low complexity domain of *SNF5*. This domain was deleted in the *ΔQsnf5* strain. Glutamine rich domains are highlighted in orange. The 4/7 histidines that were mutated to alanine in the *_HtoA_SNF5* allele are highlighted. **B)** The log_2_ of the frequency of each amino acid within QLCs divided by the global frequency of each amino acid in the proteome (*S. cerevisiae*). Values > 0 indicate enrichment in QLCs. **C)** Left: Schematic of the SWI/SNF complex engaged with a nucleosome. The *SNF5* C-termius is shown in grey, while the N-terminal QLC is shown in orange. Right: Schematic of the three main *SNF5* alleles used in this study. **D)** RT-qPCR results assessing levels of endogenous *ADH2* mRNA in four strains grown in glucose (left) or after 4 h of glucose starvation (right). Note: y-axes are different for each plot. **E)** Representative histograms (10,000 cells) showing the fluorescent signal from a *P_ADH2_-mCherry* reporter gene for four strains grown in glucose (left) or after 6 h of glucose starvation (right).

The N-terminus of *SNF5* contains two QLCs as defined above, but is overall very glutamine rich, and therefore, for simplicity, we refer to this entire 282 amino-acid region as the *SNF5* QLC from this point. We compared the sequences of Snf5 N-terminal domains taken from twenty orthologous proteins from a range of fungi (**Figure 1 – figure supplement 3**). Despite the relatively poor sequence conservation across the N-terminal disordered regions in *SNF5* (**Figure 1 – figure supplement 3A**), every region consisted of at least 18% glutamine (max 43%) and all possessed multiple histidine residues (**Figure 1 – figure supplement 3B; Supplemental Table 2**; the phylogeny considered and the total number of QLCs for each species are shown in **Figure 1 – figure supplement 3C**). In summary, enrichment for glutamine residues interspersed with histadine residues appears to be conserved sequence feature, both in QLCs in general, and in the N-terminus of SNF5 in particular, implying a possible functional role (40).

To further investigate the functional importace of the glutamine-rich N-terminal domain in *SNF5* we engineered 3 *SNF5* mutant strains: a complete deletion of the *SNF5* gene (*snf5Δ*); a deletion of the N-terminal QLC (*ΔQsnf5*); and an allele with 4 Histidines within the QLC mutated to Alanine (*_HtoA_snf5*) (Figure 1A, C).

As previously reported (34), *snf5Δ* strains grew slowly, (**Figure 1 – figure supplement 4A**). In contrast, growth rates of *ΔQsnf5* and *_HtoA_snf5* were similar to WT during continuous growth in either fermentable (glucose) or poor (galactose/ethanol) carbon sources (**Figure 1 – figure supplement 4A, B**). However, a strong growth defect was revealed for *ΔQsnf5* and *_HtoA_snf5* strains when cells were carbon starved for 24 h and then switched to a poor carbon source (Fig 1 sup 2C), suggesting that the *SNF5* QLC is important for adaptation to new carbon sources. Deletion of the *SNF5* gene has been shown to disrupt the architecture of the SWI/SNF complex leading to loss of other subunits (25, 41). To test if deletion of the QLC leads to loss of *Snf5p* protein or failure to incorporate into SWI/SNF, we immunoprecipitated the SWI/SNF complex from strains with a TAP tag at the C-terminal of the core *SNF2* subunit. We found that the entire SWI/SNF complex remained intact in both the *ΔQsnf5* and *_HtoA_snf5* strains (**Figure 1 – figure supplement 5A**). Silver-stains of the untagged *Snf5p* and Western blotting of TAP-tagged *SNF5* (42) strains showed that all *SNF5* alleles were expressed at similar levels to wild-type both in glucose and upon carbon starvation (**Figure 1 – figure supplement 5B**). Together, these results show that deletion of the *SNF5* QLC is distinct from total loss of the *SNF5 gene* and that this N-terminal sequence is important for efficient recovery from carbon starvation.

We hypothesized that slow recovery of *ΔQsnf5* and *_HtoA_snf5* strains after carbon starvation was due to a failure in transcriptional reprogramming. The alcohol dehydrogenase *ADH2* gene is normally repressed in the presence of glucose and strongly induced upon carbon starvation. This regulation depends on SWI/SNF activity (26). Therefore, we used *ADH2* as a model gene to test our hypothesis. Using reverse transcriptase quantitative polymerase chain reaction (RT-qPCR), we found that robust *ADH2* expression after acute carbon starvation was dependent on the *SNF5* QLC and the histidines within (Figure 1D). This defect was far stronger in the *ΔQsnf5* and *_HtoA_snf5* strains than in *snf5Δ* strains; *snf5Δ* strains did not completely repress *ADH2* expression in glucose, and showed partial induction upon carbon starvation, while *ΔQsnf5* strains tightly repressed *ADH2* in glucose (similar to WT), but completely failed to induce expression upon starvation (Figure 1D). These results suggest a dual-role for *SNF5* in *ADH2* regulation, both contributing to strong repression in glucose, and robust induction upon carbon starvation. The *ΔQsnf5* and *_HtoA_snf5* alleles separate these functions, maintaining WT-like repression while showing a strong defect in induction.

The RT-qPCR assay reports on the average behavior of a population. To enable single-cell analysis, we engineered a reporter strain with the mCherry (43) fluorescent protein under the control of the *ADH2* promoter integrated into the genome immediately upstream of the endogenous *ADH2* locus (Figure 1E**, Figure 1 – figure supplement 6A**). We found high cell-to-cell variation in the expression of this reporter in WT strains: after 6 h of glucose starvation, *P_ADH2_-mCherry* expression was bimodal; about half of the cells had high mCherry fluorescence and half were low. This bimodality was strongly dependent on preculture conditions, and was most apparent upon acute withdrawal of carbon from early log-phase cells (O.D. < 0.3, see methods). Complete deletion of *SNF5* eliminated this bimodal expression pattern; again, low levels of expression were apparent in glucose and induction during starvation was attenuated. As in the RT-qPCR analysis, the *ΔQsnf5* strain completely failed to induce the *P_ADH2_-mCherry* reporter at this time point and mutation of four central histidines to alanine was sufficient to mostly abrogate expression (Figure 1E). Mutation of a further two histidines had little additional effect (**Figure 1 – figure supplement 6B-D**). Taken together, these results suggest that the dual function of *SNF5* leads to switch-like control of *ADH2* expression. In glucose, *SNF5* helps repress *ADH2*. Upon carbon starvation, *SNF5* is required for efficient induction of *ADH2*. The *SNF5* QLC and histidine residues within seem to be crucial for switching between these states.

### The *SNF5* QLC is required for *ADH2* expression and recovery of neutral pH

Multiple stresses, including glucose-starvation, have been shown to cause a decrease in the pH of the cytoplasm and nucleus (nucleocytoplasm) (8, 9, 13, 44). Herein, we refer to nucleocytoplasmic pH as *intracellular pH*, or pH_i_. To investigate the relationship between *ADH2* expression and pH_i_, and how these factors depend upon *SNF5*, we engineered strains bearing both the ratiometric fluorescent pH-reporter, pHluorin (45), and the *P_ADH2_-mCherry* reporter. These cell lines allow us to simultaneously monitor pH_i_ and expression of *ADH2*.

Wild-type cells growing exponentially in 2% glucose had a pH_i_ of ∼ 7.8. Upon acute carbon starvation, cells rapidly acidified to pH_i_ ∼ 6.5. Then, during the first hour, two populations arose: an acidic population (pH_i_ ∼ 5.5), and a second population that recovered to pH_i_ ∼ 7 (Figure 2A). Cells at pH_i_ 7 proceeded to strongly induce expression of the *P_ADH2_-mCherry* reporter, while cells at pH_i_ 5.5 did not. After 8 h of glucose-starvation > 70% of wild-type cells had induced *ADH2* (Figure 2A, C).

**Figure 2:**
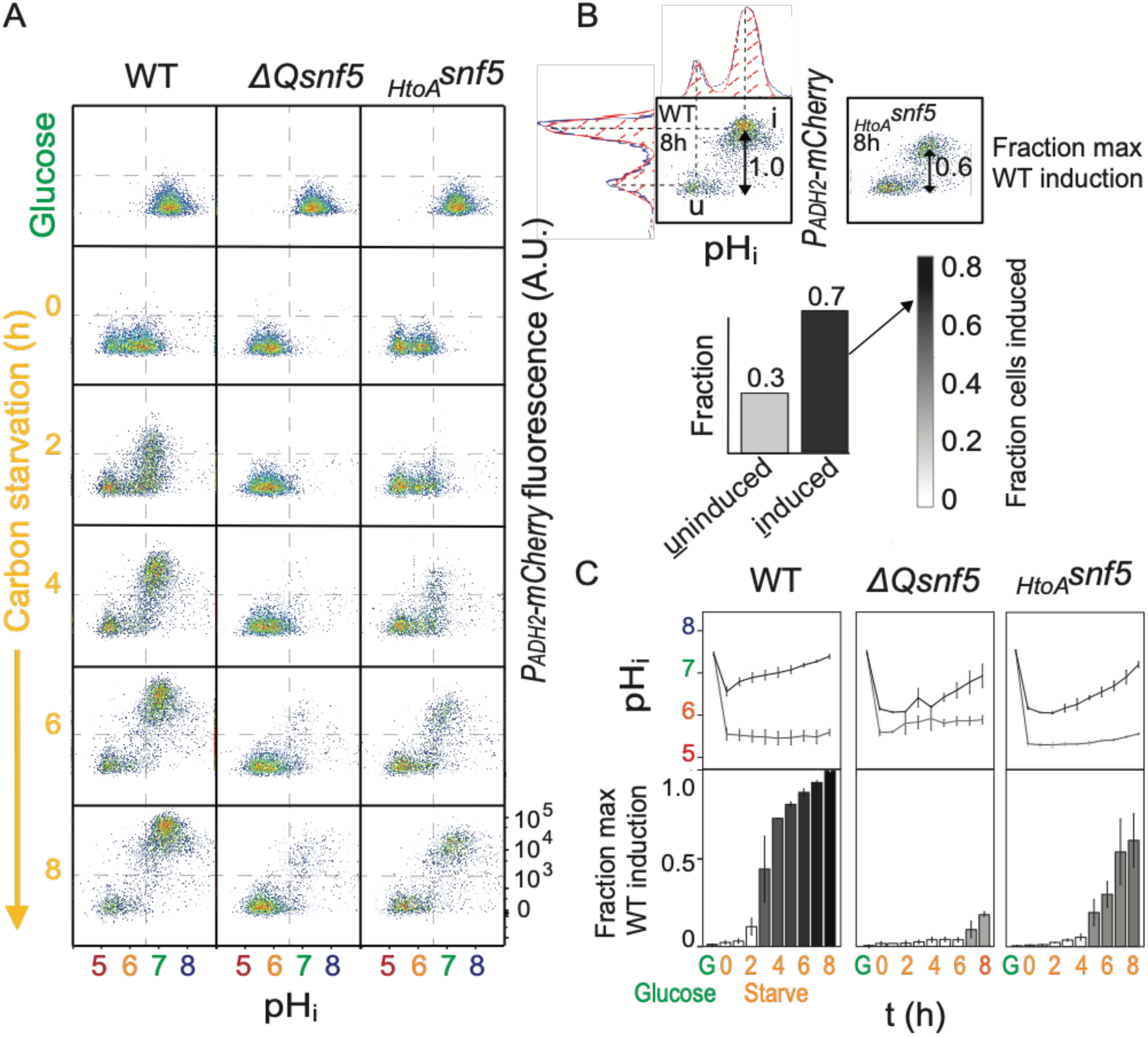
The *SNF5* QLC is required for *ADH2* expression and recovery of neutral pH. **A)** Representative flow cytometry for WT, *ΔQsnf5*, or *_HtoA_snf5* strains: the **x-axis** shows nucleocytosoplasmic pH (pH_i_), while the **y-axis** shows fluorescence from the *P_ADH2_-mCherry* reporter. Panels show cells grown in glucose (top) and then (2^nd^ to bottom) after 0 - 8 h of acute glucose-starvation. **B)** Schematic of quantification scheme: Raw data from A was fit to a single or double Gaussian curve determined by a least-residuals method. **C)** Quantification of pH_i_ and *P_ADH2_-mCherry* expression during acute starvation. The median of each Gaussian for pHi is plotted in (**C, top**). The height of bars in (**C, bottom**) indicate the fraction of maximal *P_ADH2_-mCherry* reporter gene expression (WT cells, 8 h glucose starvation) The darkness of the bars indicates the fraction of the population in the induced versus uninduced state. Mean and standard deviation of three biological replicates are shown.

We next analyzed cells harboring mutant alleles of the QLC of *SNF5*. Similarly to WT, both *ΔQsnf5* and *_HtoA_snf5* strains rapidly acidified upon carbon starvation. However, these strains were defective in subsequent neutralization of pH_i_ and in the expression of *P_ADH2_-mCherry*. At the 4 h time point, > 95 % of both *ΔQsnf5*, and *_HtoA_snf5* cells remained acidic with no detectable expression, while > 60% of wild-type cells had neutralized and expressed mCherry (Figure 2A, C). These results demonstrate that the *SNF5* QLC is necessary for efficient recovery from transient acidification. Eventually, after 24 h, the majority of mutant cells neutralized to pH_i_ ∼ 7 and induced expression of *P_ADH2_-mCherry* (**Figure 2 – figure supplement 1**). Thus, the *SNF5* QLC and histidines within are required for the rapid dynamics of both transient acidification and transcriptional induction of *P_ADH2_-mCherry* upon acute carbon starvation.

We hypothesized that mutant cells might fail to recover from acidification because transcripts controlled by SWI/SNF are responsible for pH_i_ recovery. In this model, SWI/SNF drives expression of a set of genes that must be both transcribed and translated. To test this idea we measured pH_i_ in WT cells during carbon starvation in the presence of the cyclohexamine to prevent translation of new transcripts. In these conditions, we found that cells experienced a drop in pH_i_ but were unable to recover neutral pH (**Figure 2 – figure supplement 2**). Thus, new gene expression is required for recovery of pH_i_.

### Transient acidification is required for *ADH2* induction upon carbon starvation

The acidification of the yeast nucleocytoplasm has been shown to depend upon an acidic extracellular pH (pH_e_). We took advantage of this fact to manipulate the changes in pH_i_ that occur upon carbon starvation. Cell viability was strongly dependent on pH_e_, decreasing drastically when cells were starved for glucose in media at pH ≥ 7.0 for 24 h (**Figure 3 – figure supplement 1**). Expression of *P_ADH2_-mCherry* expression was also highly dependent on pH_e_, especially in *SNF5* QLC mutants (Figure 3A**, Figure 3 – figure supplement 2**). WT cells failed to induce *P_ADH2_-mCherry* at pH_e_ ≥ 7, but induced strongly at pH_e_ ≤ 6.5. RT-qPCR showed similar behavior for the endogenous *ADH2* transcript (**Figure 3 – figure supplement 2**). Furthermore, we found that the nucleocytoplasm of all strains failed to acidify when the environment was held at pH_e_ ≥ 7 (**Figure 3 – figure supplement 3**). Therefore, we conclude that an acidic extracellular environment is required for a drop in intracellular acidity upon carbon starvation, and that this intracellular acidification is required for activation of *ADH2* transcription.

**Figure 3:**
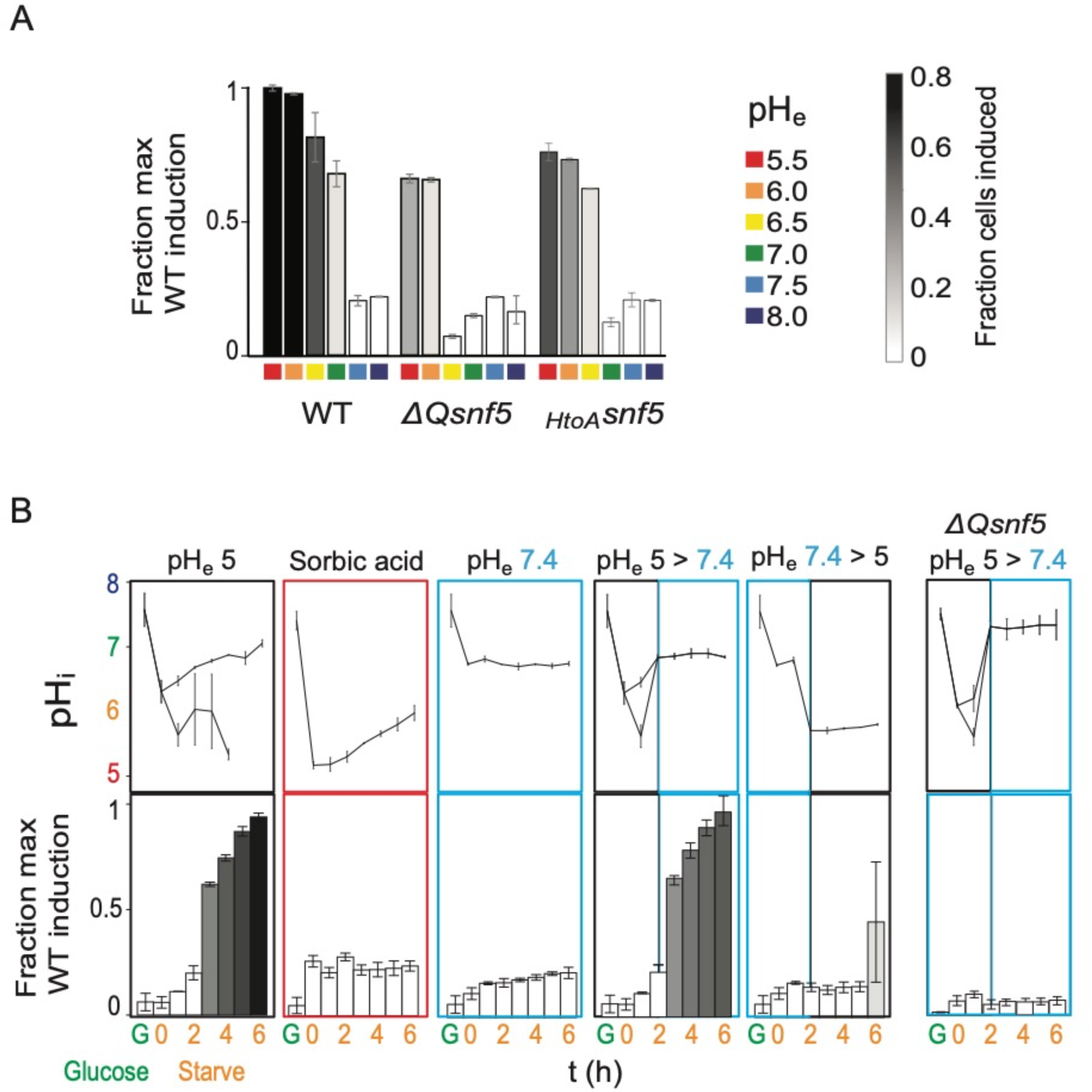
Transient acidification is required for *ADH2* induction upon carbon starvation. **A)** Expression of *P_ADH2_-mCherry* reporter gene in WT, *ΔQsnf5*, or *_HtoA_snf5* strains 8 h after acute carbon starvation in media titrated to various pH (pH_e_, see legend, right). Bar height indicates the fraction of maximal *P_ADH2_-mCherry* reporter gene expression (WT cells, pH_e_ 5.5). The darkness of the bars indicates the fraction of the population in the induced versus uninduced state (see legend, right). **B)** Time courses of glucose starvation with media manipulations to perturb the intracellular pH response, either by changing media pH (pH_e_), or by adding sorbic acid. Top panels show nucleocytoplasmic pH (pH_i_), bottom panels quantify expression of the *P_ADH2_-mCherry* reporter gene (as in A). All strains are WT except for the far right panels, which are from a *ΔQsnf5* strain.

Given that intracellular acidification is necessary for *ADH2* promoter induction, we next wondered if it was sufficient. First, we used the membrane permeable sorbic acid to allow intracellular acidification but prevent pH_i_ recovery. These cells failed to induce *P_ADH2_-mCherry*, indicating that nucleocytoplasmic acidification is not sufficient; subsequent neutralization is also required. Carbon starvation at pH_e_ 7.4 prevented transient acidification and likewise prevented expression (Figure 3B**, Figure 3 – figure supplement 3**). Cells that were first held at pH_e_ 7.4, preventing initial acidification, and then switched to pH_e_ 5, thereby causing late acidification, failed to express mCherry after 6 h. Finally, starvation at pH_e_ 5 for 2 h followed by a switch to pH_e_ 7.4, with a corresponding increase in pH_i_ led to robust *P_ADH2_-mCherry* expression. Together, these results suggest that transient acidification immediately upon switching to carbon starvation followed by recovery to neutral pH_i_ is the signal for the efficient induction of *P_ADH2_-mCherry*.

Deletion of the *SNF5* QLC leads to both failure to neutralize pH_i_ and loss *ADH2* expression. We therefore wondered if forcing cells to neutralize pH_i_ would rescue *ADH2* expression in a *ΔQsnf5* strain. This was not the case: the *ΔQsnf5* strain still fails to express *P_ADH2_-mCherry*, even if we recapitulate normal intracellular transient acidification (Figure 3B**, left**). Therefore, the *SNF5* QLC is require for normal kinetics of transient acificication *and* for additional steps in *ADH2* gene activation.

### The *SNF5* QLC and acidification of the nucleocytoplasm are required for efficient widespread transcriptional reprogramming upon carbon starvation

We wondered if transient acidification and the QLC of *SNF5* were important for transcriptional reprogramming on a genome-wide scale. To test this, we performed Illumina RNA-sequencing analysis on triplicates of each strain (WT, *ΔQsnf5*, *_HtoA_snf5*) either growing exponentially in glucose or after acute carbon-starvation for 4 h at pH_e_ 5. In addition, to test the pH-dependence of the transcriptional response, we analyzed WT strains carbon-starved at pH_e_ 7, which prevents intracellular acidification (Figure 3B**; Figure 3 – figure supplement 4**).

Principal component analysis showed tight clustering of all exponentially growing samples, indicating that mutation of the QLC of *SNF5* doesn’t strongly affect gene expression in rich media (Figure 4A). In contrast, there are greater differences between wild-type strains with mutant *SNF5* alleles upon glucose starvation. The genes that accounted for most variation (the first two principle components) were involved in carbon transport, metabolism and stress responses. We defined a set of 89 genes that were induced (> 3-fold) and 60 genes that were down-regulated (> 3-fold) in WT strains upon starvation in media titrated to pH_e_ 5. Many of these genes were poorly induced in *ΔQsnf5* and *_HtoA_snf5* mutants, as well as in WT strains starved in media titrated to suboptimal pH_e_ 7 (Figure 4B). Figures 4C and D show transcriptional differences between glucose-starved strains as volcano plots, emphasizing large-scale differences between WT and *ΔQsnf5* strains, and similarities between *ΔQsnf5* and *_HtoA_snf5*.

**Figure 4:**
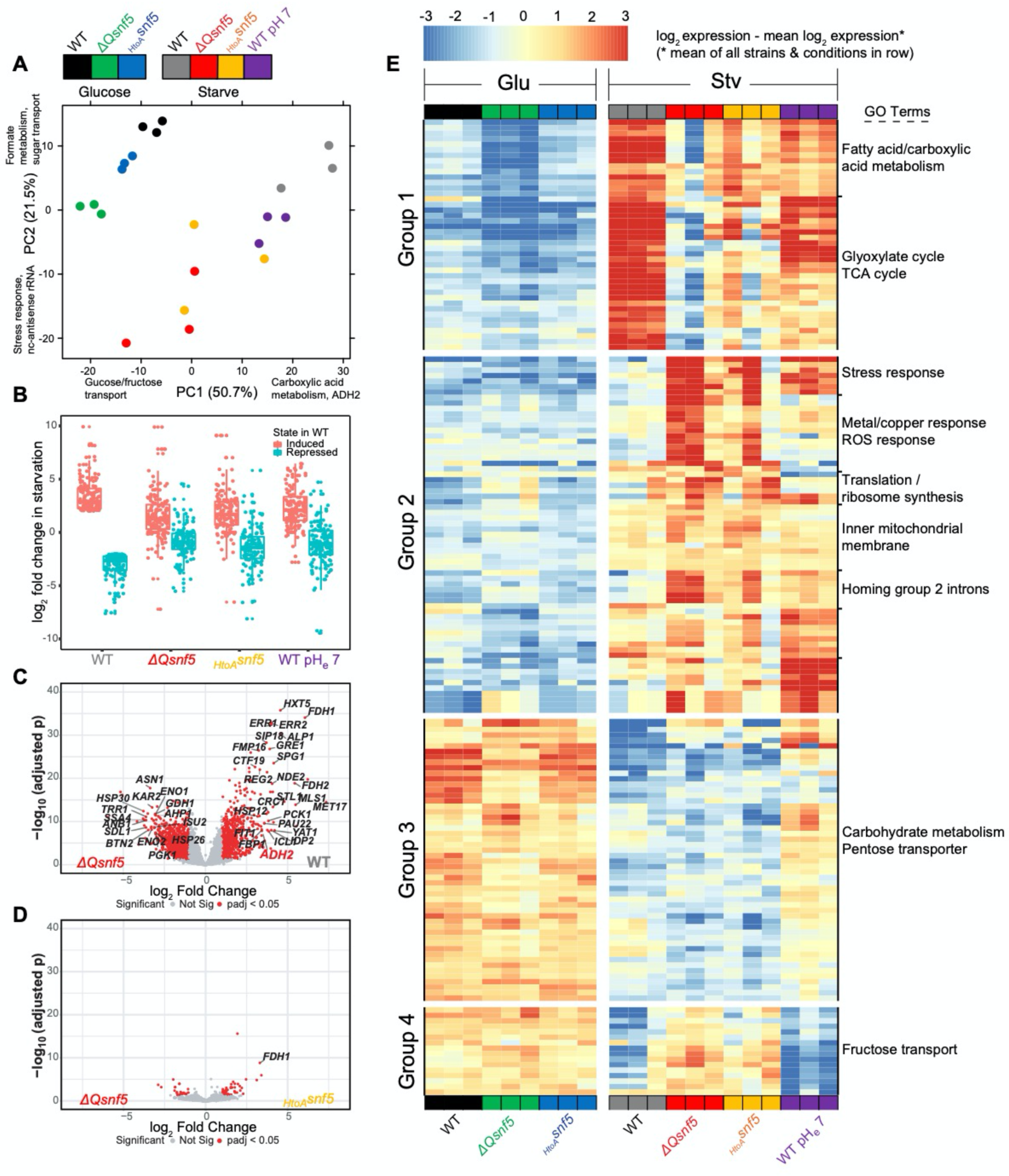
The *SNF5* QLC and acidification of the nucleocytoplasm are required for efficient widespread transcriptional reprogramming upon carbon starvation. **A)** Principal component (PC) analysis of 3 RNA-seq biological replicates for each condition tested. **B)** Expression levels of genes that were > 3 fold induced or repressed upon carbon starvation in WT strains are plotted for each *SNF5* allele. **C)** Volcano plot showing the log_2_ ratio of expression levels in WT versus *ΔQsnf5* strains (x-axis) and p-values for differential expression (y-axis). Genes with significantly different expression are indicated in red (log_2_ fold change > 1 and Wald test adjusted p value < 0.05). **D)** Volcano plot as in (C) but comparing expression levels in *_HtoA_snf5* strains to *ΔQsnf5* strains. **E)** Hierarchically clustered heat map showing expression values of 149 genes with a significant change in expression upon starvation of WT cells (log_2_ fold change > 1 and Wald test adjusted p value < 0.05). Color code indicates gene expression relative to the mean expression of that gene across all strains and conditions, with red indicating high, and blue low values (see legend). Three biological replicates are shown for each experiment. Strain and condition identities are indicated at the bottom of each column. Four groups of genes with similar behavior are indicated to the left. Gene ontology enrichment results for 9 clusters of genes are shown to the right.

We next performed hierarchical clustering analysis (Euclidean distance) of the 149 genes that are strongly differentially expressed between strains, or at suboptimal pH_e_ 7 (Figure 4E). Based on this clustering and some manual curation, we assigned these genes to four groups. Group 1 genes (n = 42) were activated in starvation in a *SNF5* QLC and pH-dependent manner. They are strongly induced in WT but induction is attenuated both in mutants of the *SNF5* QLC and when the transient acidification of pH_i_ was prevented by starving cells in media titrated to pH_e_ 7. GO analysis revealed that these genes are enriched for processes that are adaptive in carbon starvation, for example fatty acid metabolism and the TCA cycle. Group 2 (n = 64) genes were not strongly induced in WT, but were inappropriately induced during starvation in *SNF5* QLC mutants and during starvation at pH_e_ 7. GO analysis revealed that these genes are enriched for stress responses, perhaps because the failure to properly reprogram transcription leads to cellular stress. Group 3 genes (n = 51) were repressed upon carbon-starvation in a pH-dependent but *SNF5* QLC-independent manner. They were repressed in all strains, but repression failed at pH_e_ 7Finally, Group 4 genes (n = 16) were repressed in WT cells in a pH-independent manner, but failed to repress in *SNF5* QLC mutants.

We performed an analysis for the enrichment of transcription factors within the promoters of each of these gene sets using the YEASTRACT server (46). These enrichments are summarized in **Supplemental Table 2.** Top hits for Group 1 included the *CAT8* and *ADR1* transcription factors, which have previously been suggested to recruit the SWI/SNF complex to the ADH2 promoter (47).

In conclusion, both pH changes and the *SNF5* QLC are required for correct transcriptional reprogramming upon carbon starvation, but the dependencies are nuanced. Mutation of the *SNF5* QLC or prevention of nucleocytoplasmic acidification appears to trigger a stress response (Group 2 genes). Another set of genes requires pH change for their repression upon starvation, but this pH sensing is independent of *SNF5* (Group 3). A small set of genes requires the *SNF5* QLC but not pH change for repression upon starvation (Group 4). Finally, a set of genes, including many of the traditionally defined “glucose-repressed genes”, require *both* the *SNF5* QLC *and* a pH change for their induction upon carbon starvation (Group 1). For these genes, point mutation of 4 histidines in the QLC is almost as perturbative as complete deletion of the QLC. We propose that the *SNF5* QLC senses the transient acidification that occurs upon carbon starvation to elicit transcriptional activation of this gene-set. It is striking that this set is enriched for genes involved in catabolism, TCA cycle and metabolism, given that these processes are important for energetic adaptation to acute glucose-starvation.

### The *SNF5* QLC mediates a pH-sensitive transcription factor interaction *in vitro*

We reasoned that pH_i_ changes could affect the intrinsic nucleosome remodeling activity of SWI/SNF, or alternatively might impact the interactions of SWI/SNF with transcription factors. We used a fluorescence-based strategy *in vitro* to investigate these potential pH-sensing mechanisms. A center-positioned, recombinant mononucleosome was assembled on a 200 bp DNA fragment containing a “601” nucleosome positioning sequence (48) (Figure 1A). The nucleosomal substrate contained two binding sites for the Gal4 activator located upstream, and 68 base pairs of linker DNA downstream of the nucleosome. The mononucleosome contained a Cy3 fluorophore covalently attached to the distal end of the template DNA, and Cy5 was attached to the H2A C-terminal domain. The Cy3 and Cy5 fluorophores can function as a Förster Resonance Energy Transfer (FRET) pair only when the Cy3 donor and Cy5 acceptor are within an appropriate distance (see also Li and Widom, 2004). In the absence of SWI/SNF activity, the center-positioned nucleosome has a low FRET signal, but ATP-dependent mobilization of the nucleosome towards the distal DNA end leads to an increase in FRET (49–53) (Figure 5). In the absence of competitor DNA, SWI/SNF does not require an interaction with a transcription factor to be recruited to the mononucleosome and thus intrinsic nucleosome remodeling activity can be assessed independently of recruitment. In this assay, SWI/SNF complex containing ΔQsnf5p retained full nucleosome remodeling activity (Figure 5A), as well as full DNA-stimulated ATPase activity (**Figure 5 – figure supplement 1**). Furthermore, these activities were similar at pH 6.5, 7, or 7.5. Thus, we conclude that the *SNF5* QLC does not sense pH by modifying its intrinsic ATPase and nucleosome remodeling activity, at least in this *in vitro* context.

**Figure 5:**
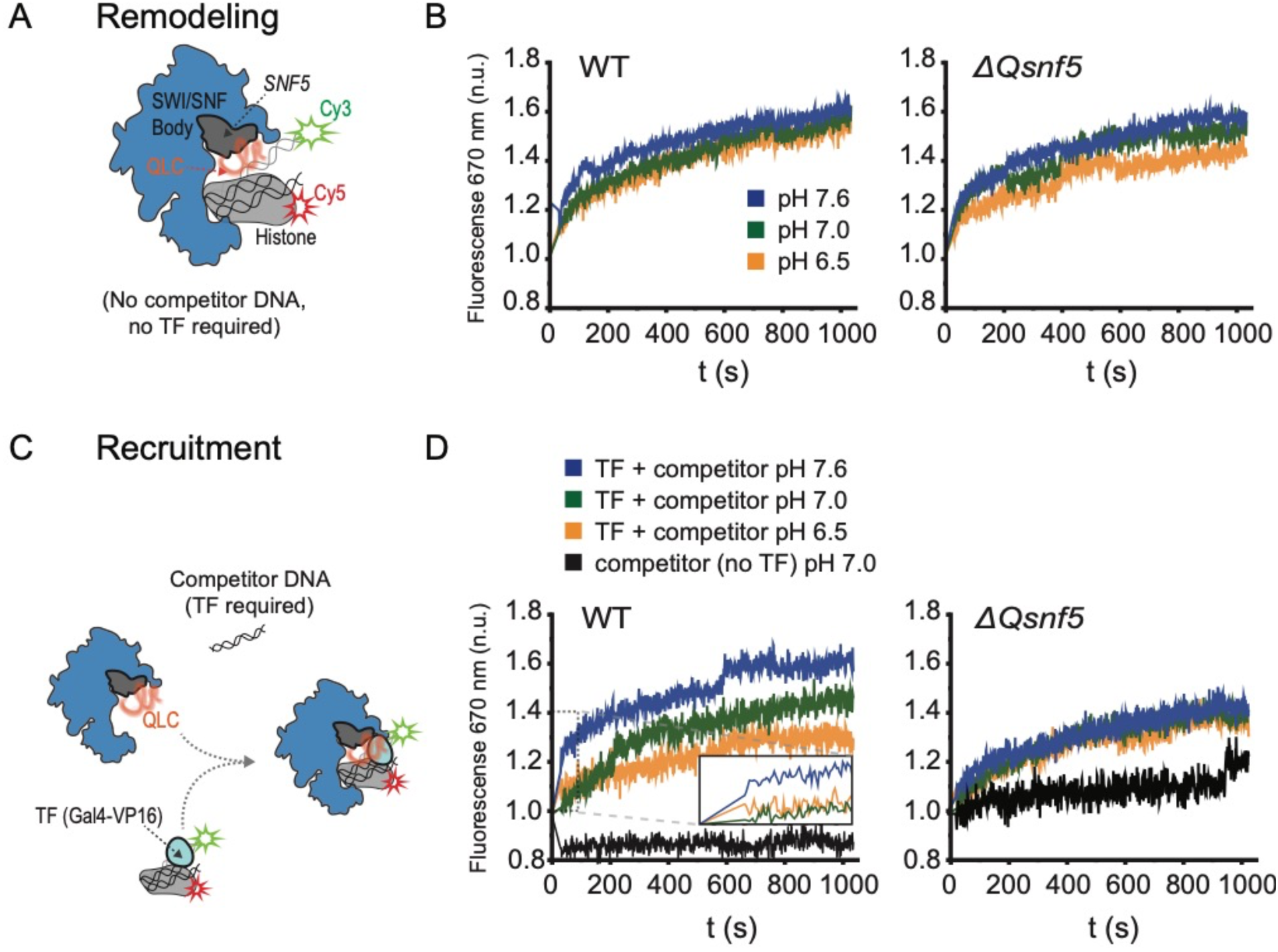
The *SNF5* QLC mediates a pH-sensitive transcription factor interaction *in vitro*. **A)** Schematic of assay: A Cy3 donor fluorophore was attached to one end of the DNA, and the histone H2A C-termini were labeled with a Cy5 acceptor fluorophore. ATP-dependent mobilization of the nucleosome to the DNA increases FRET, leading to increased emission at 670 nm. **B)** Kinetic traces for WT (left) and ΔQsnf5p (right) SWI/SNF complexes at pH 7.6 (blue), 7.0 (green), or 6.5 (orange). There is no competitor DNA, so these traces indicate intrinsic remodeling activity without requirement for recruitment by transcription factors. **C)** Schematic: In the presence of excess competitor DNA, SWI/SNF-dependent remodeling requires recruitment by a transcription factor (Gal4-VP16). **D)** Kinetic traces for WT (left) and ΔQsnf5p (right) SWI/SNF complexes at pH 7.6 (blue), 7.0 (green), or 6.5 (orange). Inset on the left panel shows the first 100 seconds of the assay after ATP addition. All traces represent FRET normalized to values prior to addition of ATP.

Next, we assessed if the *SNF5* QLC and pH changes could affect SWI/SNF interactions with transcription factors. SWI/SNF remodeling activity can be targeted to nucleosomes in vitro by Gal4 derivatives that contain acidic activation domains, an archetypal example of which is VP16 (Yudkovsky et al., 1999). Indeed, it was previously demonstrated that the QLC of Snf5p mediates interaction with the Gal4-VP16 transcription factor (32). To assess recruitment of SWI/SNF we set up reactions with an excess of nonspecific competitor DNA. In these conditions, there is very little recruitment and remodeling without interaction with a transcription factor bound to the mononucleosome DNA (Figure 5C, D). In this context, we found that the QLC of *SNF5* was required for rapid, efficient recruitment of SWI/SNF by the Gal4-VP16 activator, and that the pH of the buffer affected this recruitment (Figure 5D). Within the physiological pH-range (pH 6.5 to 7.5), recruitment and remodeling increased with pH. SWI/SNF complexes deleted for the *SNF5* QLC (containing *ΔQsnf5*p) had constitutively lower recruitment and were completely insensitive to pH changes over this same range (Figure 5D**, right**). Therefore, we conclude that the *SNF5* QLC can sense pH changes by modulating interactions between SWI/SNF and transcription factors.

### Protonation of histidines leads to conformational expansion of the *SNF5* QLC

How might pH change be sensed by *SNF5*? As described above (Figure 1B), Q-rich low-complexity sequences (QLCs) are enriched for histidines, and they are also depleted for charged amino acids (Figure 1B). Charged amino acids have repeatedly been shown to govern the conformational behavior of disordered regions (54–56). Given that histidine protonation alters the local charge density of a sequence, we hypothesized that the charge-depleted QLCs may be poised to undergo protonation-dependent changes in conformational behavior. To test this idea, we performed all-atom Monte-Carlo simulations to assess the conformational ensemble of a 50 amino acid region of the *SNF5* QLC (residues 71-120) that contained 3 histidines, 2 of which we had mutated to alanine in our experiments (Figure 6A). We performed simulations with histidines in both uncharged and protonated states to mimic possible charges of this polypeptide at the pH found in the nucleocytoplasm in glucose and carbon starvation respectively. These simulations generated ensembles of almost 50,000 distinct conformations (representative images shown in Figure 6B). To quantify conformational changes, we examined the radius of gyration, a metric that describes the global dimensions of a disordered region (Figure 6C). Protonation of the wildtype sequence led to a striking increase in the radius of gyration, driven by intramolecular electrostatic repulsions (Figure 6D**, left**). In contrast, when 2/3 histidines were replaced with alanines, no such change was observed (Figure 6D**, right**). For context, we also calculated an apparent scaling exponent (*ν^app^)*, a dimensionless parameter that can also be used to quantify chain dimensions. This analysis showed that protonation of the wildtype sequence led to a change in *ν^app^* from 0.48 to 0.55, comparable to the magnitude of changes observed in previous studies of mutations that fundamentally altered intermolecular interactions in other low-complexity disordered regions (56, 57). These results suggest that small changes in sequence charge density can elicit a relatively large change in conformational behavior. An analogous (albeit less pronounced) effect was observed for the second QLC subregion that we mutated (residues 195-233) (**Figure 6 – figure supplement 1**). Taken together, our results suggest that charge-depleted disordered regions (such as QLCs) are poised to undergo pH-dependent conformational re-arrangement. This inference offers the beginnings of a mechanism for pH-sensing by SWI/SNF: the conformational expansion of the QLC sequence upon nucleocytoplasmic acidification may tune the propensity for SWI/SNF to interact with transcription factors (Figure 6E).

**Figure 6:**
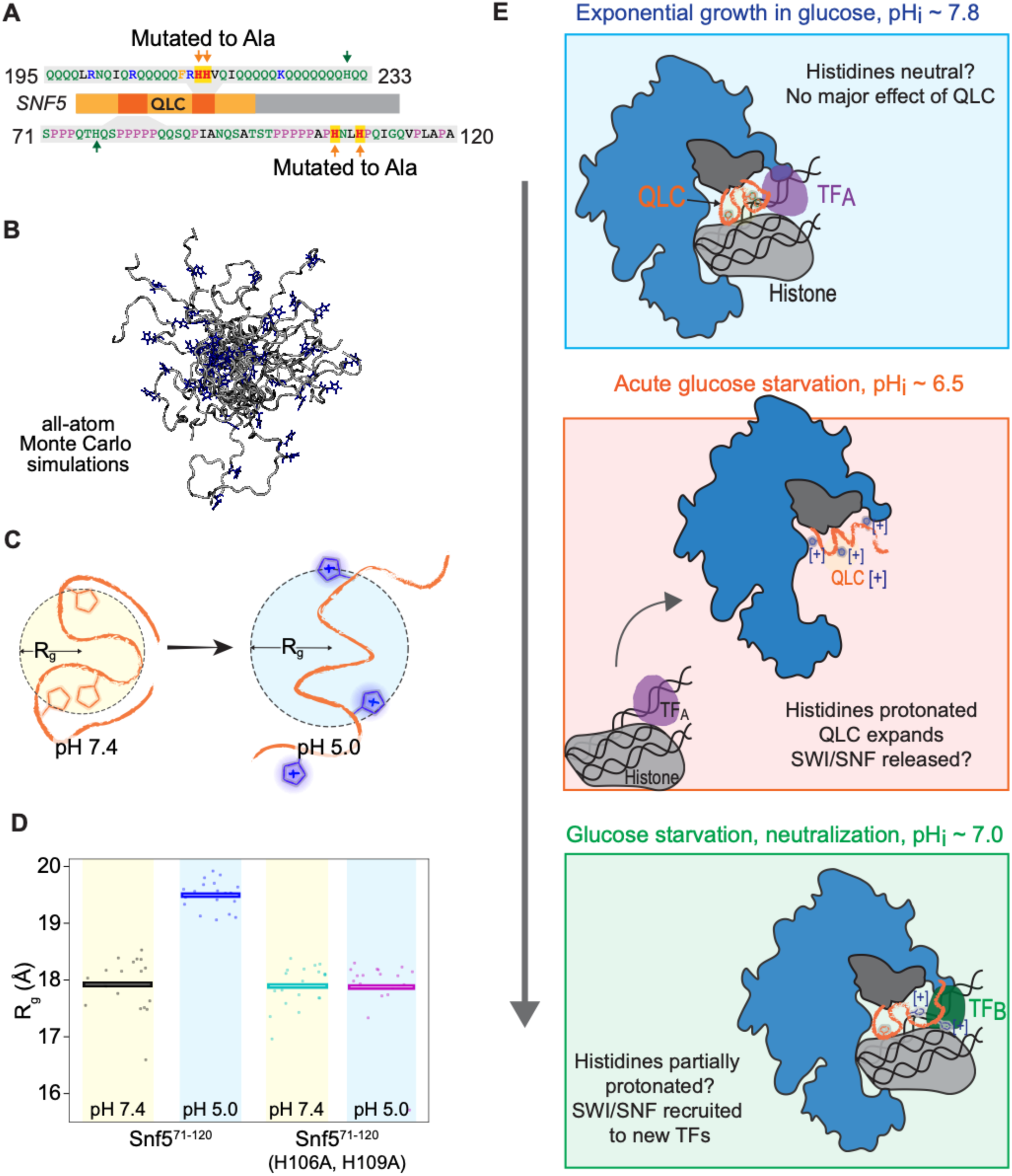
Protonation of histidines leads to conformational expansion of the *SNF5* QLC. **A)** Schematic of the *SNF5* gene (center) with the N-terminal QLC in orange, and the two simulated peptides in dark orange. Sequences of the simulated peptides and identities of histidines mutated in both the *_HtoA_snf5* yeast strain and in simulations are indicated. **B)** Representitive images of conformations sampled in Monte-Carlo all-atom simulations. **C)** Cartoon depicting quantification of radius of gyration (R_g_). **D)** Radius of gyration (**R_g_**, **y-axis**) of simulations of amino acids 71-120 of the *SNF5* QLC with histidines either neutral (pH 7.4) or protonated (pH 5.0). Left two datasets are for the native peptide, right two datasets are with 2/3 histidines (H106 and H109) replaced with alanine, mimicking the *_HtoA_snf5* allele. Points represent the mean R_g_ from all conformations sampled in each independent simulation (beginning from distinct random initial conformers). Bars represent the mean values of all simulations. **E)** Model of SWI/SNF regulation during carbon starvation. Top) In glucose (pH_i_ ∼ 7.8), the *SNF5* QLC is unprotonated. SWI/SNF is engaged by transcription factors that prevent transcription of glucose repressed genes, or that activate other genes (TF_A_). Middle) Upon acute carbon starvation, pH_i_ drops to ∼ 6.5 leading to protonation of histidines in the *SNF5* QLC. Conformational expansion of the QLC may aid the release of SWI/SNF from some transcription factors (TF_A_), and potentially drive recruitment to others (not shown). Bottom) As the cell adapts to carbon starvation, pH_i_ neutralizes to ∼ 7.0. Histidines within the *SNF5* QLC may be partially protonated? The pK_a_ of histidine is highly context-dependent. The QLC may aid recruitment of SWI/SNF to the promoters of glucose-repressed genes, thus leading to their expression.

## Discussion

Intracellular pH changes occur in many physiological contexts, including cell cycle progression (58), the circadian rhythm of crassulacean acid metabolism plants (59), oxidative stress (60), heat shock (13), osmotic stress, (61), and changes in nutritional state (15, 62). However, the physiological role of these pH_i_ fluctuations, and the molecular mechanisms to detect them, remain poorly understood. Prior results have emphasized the inactivation of processes in response to cytosolic acidification (17–19). However, it is unclear how necessary modifications to the cell can occur if cellular dynamics are uniformly decreased. Much less has been reported regarding a potential role of fluctuations in pH_i_ as a signal to activate specific cellular programs. In this work, we found that transient acidification is required for activation of glucose-repressed genes. Therefore, our work establishes a positive regulatory role for nucleocytoplasmic pH changes during carbon starvation.

Previous studies of intracellular state during glucose starvation based on population averages reported a simple decrease in pH_i_ (15). In this work, we used single-cell measurements of both pH_i_ and gene expression, and found that two co-existing subpopulations arose upon acute glucose-starvation, one with pH_i_ ∼ 5.5 and a second at ∼ 6.5. The latter population recovered to neutral pH_i_ and then induced glucose-repressed genes, while the former remained dormant in an acidified state. We have not yet determined the mechanism that drives the bifurcation in pH response. It is possible that this bistability provides a form of bet-hedging (63) where some cells attempt to respond to carbon starvation, while others enter a dormant state (19). However, we have yet to discover any condition where the population with lower pH_i_ and delayed transcriptional activation has an advantage. An alternative explanation is that these cells are failing to correctly adapt to starvation, perhaps undergoing a metabolic crisis, as suggested in a recent study (62).

It is becoming clear that intracellular pH is an important mechanism of biological control. It was previously shown that the protonation state of phosphatidic acid (PA) determines binding to the transcription factor Opi1, coupling membrane biogenesis and intracellular pH (4). We focused our studies on the N-terminal region of *SNF5* because it is known to be important for the response to carbon starvation and contains a large low-complexity region enriched in both glutamine and histidine residues. Histidines are good candidates for pH sensors as they can change protonation state over the recorded range of physiological pH fluctuations, and their pK_a_ can be tuned substantially depending on local sequence context. Consistent with this hypothesis, we found that the *SNF5* QLC and the histidines embedded within were required for transcriptional reprogramming.

Global analysis revealed that genes that require pH_i_ oscillation and the *SNF5* QLC for their induction during carbon starvation are involved in metabolic processes including the TCA cycle, fatty acid metabolism and the glyoxylate cycle. The upregulation of these metabolic pathways may provide alternative energy sources. It will be interesting to see if human SWI/SNF undergoes similar pH-dependent regulation. Cancer biology hints that this may be the case. It has been observed that about 20% of human cancers have mutations in the SWI/SNF complex (64). Human *SNF5* (SMARCB1) was the first subunit of the SWI/SNF to be linked to cancer, where it is mutated in most cases of pediatric malignant rhabdoid tumor (65, 66). It is known that mutations of the SWI/SNF that lead to cancer generally result in misregulation of fatty acid synthesis, which is required for cancer proliferation (67, 68). The pH-sensing QLC found in yeast *SNF5* is absent in the human orthologue, SMARCB1, but QLCs and regions of extreme histidine enrichment are present in the Arid1a, Arid1b and Arid2 subunits of human SWI/SNF, and loss of Arid1a is a leading cause of ovarian and uterine cancers (69). An acidic pH is a prominent feature of the tumor microenvironment (70, 71) and intracellular pH tends to be elevated in tumor cells. These observations motivate the future study of pH-sensing by SWI/SNF in humans.

Our *in vitro* assays showed that the intrinsic ATPase and nucleosome remodeling activities of SWI/SNF are robust to pH changes from 6.5 to 7.6. However, recruitment of remodeling activity by a model transcription factor (GAL4-VP16) was pH-sensitive, and this pH dependence was dependent on the *SNF5* QLC. In this case, the recruitment by GAL4-VP16 was inhibited at pH 6.5. We speculate that low pH_i_ favors release of SWI/SNF from activators that it is bound to in glucose conditions, and then the subsequent partial recovery in pH_i_ could allow it to bind to a different set of activators, thus recruiting it to genes that are expressed during starvation. This model is consistent with the requirement for both acidification and subsequent neutralization for expression of *ADH2* (Figure 3). In principle, the conformational dynamics of the *SNF5* QLC could be distinct at all three stages (Figure 6E). There are almost certainly additional pH-sensing elements of the transcriptional machinery that also take part in this reprogramming.

Low complexity sequences, including QLCs, tend to be intrinsically disordered and therefore highly solvent exposed. A recent large-scale study of intrinsically disordered sequences showed that their conformational behavior is inherently sensitive to changes in their solution environment (36, 37). Similarly, our simulations revealed that histidine protonation may lead the *SNF5* QLC to expand dramatically. This provides a potential mechanism for pH-sensing: upon acidification, histidines become positively charged leading QLCs to adopt a more expanded state, perhaps revealing short linear interaction motifs (SLIMs), reducing the entropic cost of binding to interaction partners, preventing polar-mediated protein-protein interactions, or facilitating electrostatic mediated contacts. The enrichment of histidines in QLCs hints that this could be a general, widespread mechanism to regulate cell biology in response to pH changes.

Glutamine-rich low-complexity sequences have been predominantly studied in the context of disease. Nine neurodegenerative illnesses, including Huntington’s disease, are thought to be caused by neurotoxic aggregation seeded by proteins that contain polyglutamines created by expansion of CAG trinucleotide repeats (72). However, polyglutamines and glutamine-rich sequences are relatively abundant in *Eukaryotic* cells: More than 100 human proteins contain QLCs, and the *Dictyostelium* and *Drosophilid* phyla have QLCs in ∼ 10% and ∼ 5% of their proteins respectively (73). Furthermore, there is clear evidence of purifying selection to maintain polyQs in the *Drosophilids* (74). This prevalence and conservation suggest an important biological function for these sequences. Recent work in *Ashbya gosypii* has revealed a role for QLC-containing proteins in the organization of the cytoplasm through phase separation into liquid droplets to enable subcellular localization of signaling molecules (75). More generally, polyglutamine has been shown to drive self-association into a variety of higher-order assemblies, from fibrils to nanoscopic spheres to liquid droplets (76–78). Taken together, these results imply that QLCs may offer a general mechanism to drive protein-protein interactions. In this study, we have identified a role for QLCs in the SWI/SNF complex as pH-sensors. Our current model (Figure 6E) is that the *SNF5* QLC partakes in heterotypic protein interactions that are modulated by protonation of histidines when the cell interior acidifies. However, we don’t rule out the possibility for homotypic interactions and higher-order assembly of multiple SWI/SNF complexes.

All cells must modify gene expression to respond to environmental changes. This phenotypic plasticity is essential to all life, from single celled organisms fighting to thrive in an ever-changing environment, to the complex genomic reprogramming that must occur during development and tissue homeostasis in plants and *metazoa*. Despite the differences between these organisms, the mechanisms that regulate gene expression are highly conserved. Changes in intracellular pH are increasingly emerging as a signal through which life perceives and reacts to its environment. This work provides a new role for glutamine-rich low-complexity sequences as molecular sensors for these pH changes.

## Material and Methods

### Cloning and yeast transformations

Yeast strains used in this study were all in the S288c strain-background (derived from BY4743). The sequences of all genes in this study were obtained from the *Saccharomyces cerevisiae* genome database (http://www.yeastgenome.org/).

We cloned the various *SNF5* alleles into plasmids from the Longtine/Pringle collection (79). We assembled plasmids by PCR or gene synthesis (IDT gene-blocks) followed by Gibson cloning (80). Then, plasmids were linearized and used to overwrite the endogenous locus by sigma homologous recombination using homology to both ends of the target gene.

The *ΔQsnf5* gene lacks the N-terminal 282 amino acids that comprise a glutamine rich low complexity domain. Methionine 283 serves as the ATG for the *ΔQ-SNF5* gene. In the *_HtoA_snf5* allele, histidines 106, 109, 213 and 214 were replaced by alanine using mutagenic primers to amplify three fragments of the QLC region which were combined by Gibson assembly into a *SNF5* parent plasmid linearized with BamH1 and Sac1.

We noticed that the slow growth null strain phenotype of the *snf5Δ* was partially lost over time, presumably due to suppressor mutations. Therefore, to avoid these spontaneous suppressors, we first introduced a CEN/ARS plasmid carrying the *SNF5* gene under its own promoter and the *URA3* auxotrophic selection marker. Then a kanMX6 resistance cassette, amplified with primers with homology at the 5’ and 3’ of the *SNF5* gene was used to delete the entire chromosomal *SNF5* ORF by homologous recombination. We subsequently cured strains of the CEN/ARS plasmid carrying WT *SNF5* by negative selection against its URA3 locus by streaking for single colonies on 5-FOA plates immediately before each experiment to analyze the *snf5Δ* phenotype.

The *P_ADH2_-mCherry* reporter was cloned into integrating pRS collection plasmids (81). *URA3* (pRS306) or *LEU2* (pRS305) were used as auxotrophic selection markers. The 835 base pairs upstream of the *ADH2* gene was used as the promoter (*P_ADH2_*). *P_ADH2_*, and the mCherry ORF were amplified by PCR and assembled into linearized pRS plasmids (Sac1/Asc1) by Gibson assembly. These plasmids were cut in the middle of the *ADH2* promoter using the Sph1 restriction endonuclease and integrated into the endogenous *ADH2* locus by homologous recombination.

The *pHluorin* gene was also cloned into integrating pRS collection plasmids. *URA3* (pRS306) and *LEU2* (pRS305) were used for selection. The plasmid with the *pHluorin* gene was obtained described in (15). We amplified the *pHluorin* gene and the strong *TDH3* promoter and used Gibson assembly to clone these fragments into pRS plasmids linearized with Sac1 and Asc1. Another strategy was to clone the *pHluorin* gene and a natMX6 cassette into the integrating pRS304 plasmid (that contains *TRP1*), which was then linearized within the *TRP1* cassette using HindIII and integrated into the *TRP1* locus.

A C-terminal TAP tag was used to visualize Snf5 and Snf2 proteins in Western blots. pRS plasmids were used but the cloning strategy was slightly different. A 3’ fragment of the *SNF5* and SNF2 genes were PCR amplified without the Stop codon. This segment does not contain a promoter or an ATG codon for translation initiation. The TAP tag was then amplified by PCR and cloned together with the 3’ of *SNF5* and *SNF2* ORFs by Gibson assembly into pRS plasmids with linearized Sac1 and Asc1. Plasmids were linearized in the 3’ of the *SNF5* or *SNF2* ORFs with StuI and XbaI respectively to linearize the plasmid allowing integration it into the 3’ of each gene locus by homologous recombination. Therefore, transformation results in a functional promoter at the endogenous locus fused to the TAP tag.

The *SNF5*-*GFP* strain was obtained from the yeast GFP collection (82), a gift of the Drubin/Barnes laboratory at UC Berkeley. The *SNF2*-*GFP* fused strain was made by the same strategy used for the TAP tagged strain above.

**Supplemental Tables 6 and 7** list strains and plasmids generated in this study.

### Culture media

Most experiments, unless indicated, were performed in synthetic complete (SC) media (13.4 g/L yeast nitrogen base and ammonium sulfate; 2 g/L amino acid mix and 2% glucose). Carbon starvation media was SC media without glucose, supplemented with sorbitol, a non-fermentable carbon source to avoid osmotic shock during glucose-starvation (6.7 g/L YNB + ammonium sulfate; 2g/L Amino acid mix and 100 mM sorbitol). The pH of starvation media (pH_e_) was adjusted using NaOH.

### Glucose-starvation

Cultures were incubated in a rotating incubator at 30^Ο^C and grown overnight (14 - 16 h) to an OD between 0.2 and 0.3. Note: it is extremely important to prevent culture OD from exceeding 0.3, and results are different if cells are allowed to saturate and then diluted back. Thus, it is imperative to grow cultures from colonies on plates for > 16 h without ever exceeding OD 0.3 to obtain reproducible results. Typically, we would inoculate 3 ml cultures and make a series of 4 - 5 1/5 dilutions of this starting culture to be sure to catch an appropriate culture the following day. 3 milliliters of OD 0.2 - 0.3 culture were centrifuged at 6000 RPM for 3 minutes and re-suspended in 3 ml starvation media (SC sorbitol at various pH_e_). This spin and resuspension was repeated two more times to ensure complete removal of glucose. Finally, cells were re-suspended in 3 milliliters of starvation media. For flow cytometry, 200 μL samples were transferred to a well of a 96-well plate at each time point. During the course of time lapse experiments, culture aliquots were set aside at 4°C. The LSR II flow cytometer with the HTS automated sampler were used for all measurements. 10,000 cells were analysed at each time point.

### Nucleocytosoplasmic pH measurements

Nucleocytoplasmic pH (pH_i_) was measured by flow cytometry or microscopy. The ratiometric, pH-sensitive GFP variant, *pHluorin*, was used to measure pH based on the ratio of fluorescence from two excitation wavelengths. The settings used on our for LSR II flow cytometer were AmCyan (excitation 457, emission 491) and FITC (excitation 494, emission 520). AmCyan emission increases with pH, while FITC emission decreases. A calibration curve was made for each strain in each experiment. To generate a calibration curve, glycolysis and respiration were poisoned using 2-deoxyglucose and azide. This treatment leads to a complete loss of cellular ATP, and the nucleocytoplasmic pH equilibrates to the extracellular pH. We used the calibration buffers published by Patricia Kane’s group (83): 50 mM MES (2-(N-morpholino) ethanesulfonic acid), 50 mM HEPES (4-(2-hydroxyethyl)-1-piperazineethanesulfonic acid, 50 mM KCL, 50 mM NaCL, 0.2 M ammounium acetate, 10 mM sodium azide, 10 mM 2-Deoxyglucose. Buffers were titrated to the desired pH with HCL or NaOH. Sodium azide and 2-deoxyglucose were always added fresh.

### RT-qPCR

For qPCR and RNA seq, RNA was extracted with the “High pure RNA isolation kit” (Roche) following the manufacturer’s instructions. Three biological replicates were performed. cDNAs and qPCR were made with iSCRIPT and iTAQ universal SYBR green supermix by Bio-Rad, following the manufacturer’s instructions. Samples processed were: exponentially growing culture (+Glu), or acute glucose-starvation for 4 h in media titrated to pH 5.5 or 7.5. Primers for qPCR were taken from Biddick et al 2008; for *ADH2* and *FBP1* genes: forward (GTC TAT CTC CAT TGT CGG CTC), reverse (GCC CTT CTC CAT CTT TTC GTA), and forward (CTT TCT CGG CTA GGT ATG TTG G), reverse (ACC TCA GTT TTC CGT TGG G). *ACT1* was used as an internal control; primers were: forward (TGG ATT CCG GTG ATG GTG TT), reverse (TCA AAA TGG CGT GAG GTA GAG A).

### RNA sequencing

We performed RNA sequencing analysis to determine the extent of the requirement for the *SNF5* QLC in the activation of glucose-repressed genes. Three biological replicates were performed. Total RNA was extracted from WT, *ΔQ-snf5* and *_HtoA_snf5* strains during exponential growth (+Glu) and after 4 hours of acute glucose starvation. In addition, WT strains were acutely starved in media titrated to pH 7. Next, poly-A selection was performed using Dynabeads and libraries were performed following manufactures indications. Sequencing of the 32 samples was performed on an Illumina Hi-seq on two lanes. RNA-seq data were aligned to the University of California, Santa Cruz (UCSC), sacCer2 genome using Kallisto (0.43.0, http://www.nature.com/nbt/journal/v34/n5/full/nbt.3519.html) and downstream visualization and analysis was in R (3.2.2). Differential gene expression analysis, heat maps and volcano plots were created using DESeq2 where a Wald test was used to determine differentially expressed genes and Euclidean distance to calculate clustering for heat maps.

RNA-seq R-code can be found at: https://github.com/gbritt/SWI_SNF_pH_Sensor_RNASeq

### Western blots

Strains containing *SNF5* and *SNF2* fused to the TAP tag were used. Given the low concentration of these proteins, they were extracted with Trichloroacetic acid (TCA): 3 mL culture was pelleted by centrifugation for 2 min at 6000 RPM and then frozen in liquid nitrogen. Pellets were thawed on ice and re-suspended in 200 uL of 20% TCA, ∼ 0.4 g of glass beads were added to each tube. Samples were lysed by bead beating 4 times for 2 min with 2 min of resting in ice in each cycle. Supernatants were extracted using a total of 1 mL of 5% TCA and precipitated for 20 min at 14000 RPM at 4 C. Finally, pellets were re-suspended in 212 uL of Laemmli sample buffer and pH adjusted with ∼26 uL of Tris buffer pH 8. Samples were run on 7 - 12% gradient polyacrylamide gels with Thermo-Fisher PageRuler prestained protein ladder 10 to 18 KDa. Proteins were transferred to a nitrocellulose membrane, which was then blocked with 5% nonfat milk and incubated with a rabbit IgG primary antibody (which binds to the protein A moiety of the TAP tag) for 1 hour and then with fluorescently labelled goat anti-rabbit secondary antibody IRdye 680RD goat-anti-rabbit (LI-COR Biosciences Cat# 926–68071, 1:15,000 dilution). Anti-glucokinase was used as a loading control (rabbit-anti-Hxk1, US Biological Cat# H2035-01, RRID:AB_2629457, Salem, MA, 1:3,000 dilution) followed by IRDye 800CW goat-anti-rabbit (LI-COR Biosciences Cat# 926-32211, 1:15,000 dliution). Membranes were visualized using a LI-COR Odyssey CLx scanner with Image Studio 3.1 software. Fluorescence emission was quantified at 700 and 800 nM.

### Co-immunoprecipitation of SWI/SNF complex

For each purification, 6 L of cells were grown in YPD to an OD of 1.2. Cells were broken open using glass beads in buffer A (40 mM HEPES [K+], pH 7.5, 10% glycerol, 350 mM KCl, 0.1 % Tween-20, supplemented with 20 µg/mL leupeptin, 20 µg/mL pepstatin, 1µg/mL benzamidine hydrochloride and 100 µM PMSF) using a Biospec bead beater followed by treatment with 75 units of benzonase for 20 minutes (to digest nucleic acids). Heparin was added to a final concentration of 10 µg/mL. The extract was clarified by first spinning at 15,000 RPM in a SS34 Sorvall rotor for 30 minutes at 4°C, followed by centrifugation at 45,000 RPM for 1.5 hours at 4°C in a Beckman ultracentrifuge. The soluble extract was incubated with IgG sepharose beads for 4 hours at 4°C using gentle rotation. IgG sepharose bound proteins were washed 5 times in buffer A and once in buffer B (10 mM TRIS-HCl, pH 8.0, 10% glycerol, 150 mM NaCl, 0.5 mM EDTA, 0.1% NP40, 1 mM DTT, supplemented with 20 µg/mL leupeptin, 20 µg/mL pepstatin, 1µg/mL benzamidine hydrochloride and 100 µM PMSF). Bound protein complexes were incubated in buffer B with TEV protease overnight at 4°C using gentle rotation. The eluted protein was collected, CaCl2 was added to a final concentration of 2 mM and bound to calmodulin-sepharose beads for 4 hours at 4°C using gentle rotation. Following binding the protein-bound calmodulin-sepharose beads were washed 5 times in buffer C (10 mM TRIS-HCl, pH 8.0, 10% glycerol, 150 mM KCl, 2 mM CaCl2, 0.1% NP40, 1 mM DTT, supplemented with 20 µg/mL leupeptin, 20 µg/mL pepstatin, 1µg/mL benzamidine hydrochloride and 100 µM PMSF). The bound proteins were eluted in buffer D (10 mM TRIS-HCl, pH 8.0, 10% glycerol, 150 mM KCl, 2 mM EGTA, 0.1% NP40, 0.5 mM DTT, supplemented with 20 µg/mL leupeptin, 20 µg/mL pepstatin, 1µg/mL benzamidine hydrochloride and 100 µM PMSF. The protein complexes were resolved by SDS-PAGE and visualized by silver staining.

### Data fitting

Fluorescence intensity from the *P_ADH2_-mCherry* reporter and ratiometric fluorescence measurements from pHluorin were fit with a single or double Gaussian curve for statistical analysis using MATLAB (MathWorks). The choice of a single or double Gaussian fit was determined by assessing which fit gave the least residuals. For simplicity, the height (mode) of each Gaussian peak was used to determine the fraction of cells in each population rather than the area, because peaks overlapped in many conditions.

### Sequence analysis of QLCs

A glutamine-rich low-complexity sequence was defined as a sequence containing at least ten glutamines, within which we allowed any number of single or double amino acid insertions, but terminated by any interruption of three or more non-glutamine residues. For example, QQQQQAAQQQQQ and QAQAQAQAQAQAQAQAQAQ both count as a continuous QLCs, but QQQQQAAAQQQQQ does not. *Saccharomyces cerevisiae* genome and protein sequences (S288c) were downloaded from SGD (www.yeastgenome.org). Amino acid enrichment scores within QLCs compared to the global frequencies of amino acids in each proteome were calculated for *Saccharomyces cerevisiae*, *Drosophila melanogaster*, *Homo sapiens* and *Dictyostelium discoideum* reference protein sequences (downloaded from http://www.ebi.ac.uk) (84).

### Nucleosome Remodeling assays

#### SWI/SNF purification

SWI/SNF complexes were purified from yeast strains with a tandem affinity purification protocol as previously described (Smith et al., 2005). Cells were grown in YPAD media and harvested at OD_600_ = 3, and flash frozen and stored at −80°C. Yeast cells were lysed using a cryomill (PM100 Retsch). Ground cell powder was resuspended in E Buffer (20mM Hepes, 350mM NaCl, 0.1% Tween-20, 10% glycerol, pH 7.5), with fresh 1mM DTT and protease inhibitors (0.1 mg/mL phenylmethylsulfonyl fluoride, 2ug/mL leupeptin, 2ug/mL pepstatin, 1mM benzamidine) and incubated on ice for 30 minutes. The crude lysate was clarified first by centrifugation 3K rpm for 15 minutes, and then 40K rpm for 60 minutes at 4°C. The clear lysate was transferred to a 250 mL falcon tube and incubated with 400 uL IgG resin slurry (washed previously with E buffer without protease inhibitors) for 2 hours at 4° C. The resin was washed extensively with E buffer and protease inhibitors, and the protein-bound resin was incubated with 300 units TEV protease overnight at 4°C. The eluent was collected, incubated with 400 uL Calmodulin affinity resin, washed previously with E buffer with fresh protease inhibitors, DTT and 2mM CaCl_2_, for 2 hours at 4°C. Resin washed with the same buffer and SWI/SNF was eluted with E buffer with protease inhibitors, DTT, and 10 mM EGTA. The eluent was dialyzed in E buffer with PMSF, DTT, and 50 uM ZnCl_2_ at least 3 times. The dialyzed protein was concentrated with a Vivaspin column, aliquoted, flash frozen, and kept at −80°C. SWI/SNF concentration was quantified by electrophoresis on 10% SDS-PAGE gel alongside a BSA standard titration, followed by SYPRO Ruby (Thermo Fisher Scientific) staining overnight and using ImageQuant 1D gel analysis.

#### Mononucleosome reconstitutions

Recombinant octamers were reconstructed from isolated histones as described previously (Luger et al., 1999). In summary, recombinant human H2A (K125C), H2B, and H3 histones and *Xenopus laevis* H4 were isolated from *Escherichia coli* (Rosetta 2 (DE3) with and without pLysS). In order to label human H2A, a cysteine mutation was introduced at residue K125 via site-directed mutagenesis, which was labeled with Cy5 fluorophore attached to maleimide group (Zhou and Narlikar, 2016). DNA fragments were generated from 601 nucleosome positioning sequence and 2x Gal4 recognition sites with primers purchased from IDT. For FRET experiments, PCR amplification of labeled DNA fragments were as followed: 500nM Cy3 labeled (5’-Cy3/TCCCCAGTCACGACGTTGTAAAAC-3’) and unlabeled primers (5’-ACCATGATTACGCCAAGCTTCGG-3’), 200uM dNTPs, 0.1ng/ul p159-2xGal4 plasmid kindly donated by Blaine Bartholomew, 0.02 U/ul NEB Phusion DNA Polymerase, 1x Phusion High Fidelity Buffer. For ATPase assays, two unlabeled primers used (PrimerW: 5’-GTACCCGGGGATCCTCTAGAGTG-3’, PrimerS: 5’-GATCCTAATGACCAAGGAAAGCA-3’) under same PCR conditions with NEB Taq DNA Polymerase with 1x NEB ThermoPol Buffer. 400 nM fluorescently-labeled and unlabeled mononucleosomes were reconstituted via salt gradient at 4°C with a peristaltic pump as described previously (Luger et al., 1999), with 600mL high salt buffer (10 mM Tris-HCl, pH = 7.4, 1 mM EDTA, 2M KCl, 1 mM DTT) exchanged with 3 L of low salt buffer (10 mM Tris-HCl, pH = 7.4, 1 mM EDTA, 50 mM KCl, 1 mM DTT) over 20 hr. The quality of the nucleosomes was checked by visualizing on a 5% native-PAGE gel and scanning fluorescence ratios on ISS PC1 spectrofluorometer.

#### FRET-based nucleosome remodeling

The fluorescence resonance energy transfer between Cy3-labeled DNA and Cy5 labeled octamer is used to measure the remodeling and recruitment activity of SWI/SNF, using an ISS PC1 spectrofluorometer. The remodeling activity was measured by increase in FRET signal in response to sliding of octamer on the DNA template. The reaction was performed under three different pH conditions pH 6.5 (25 mM MES, 0.2 mM EDTA, 5 mM MgCl_2_, 70 mM KCl, 1 mM DTT), pH 7 (25 mM Tris, 0.2 mM EDTA, 5 mM MgCl_2_, 70 mM KCl, 1 mM DTT) or pH 7.6 (25 mM HEPES, 0.2 mM EDTA, 5 mM MgCl_2_, 70 mM KCl, 1 mM DTT). A remodeling reaction contained 2 nM or 4 nM (WT or mutant) SWI/SNF, 5 nM nucleosome and 100 uM ATP or AMP-PNP. A 100 seconds of pre-scan of the reaction is taken before the reaction started and the time-dependent fluorescence measurements started after addition of ATP or AMP-PNP for 1000 seconds at room temperature. Similarly, recruitment assays were performed in three different buffer conditions: pH 6.5, pH 7 and pH 7.6. The recruitment assays contained 2 nM or 4 nM (WT or mutant) SWI/SNF, 5 nM nucleosome, 4 nM competitor DNA, 100 uM Gal4–VP16 (Protein One, P1019-02) and 100 uM ATP or AMP-PNP, together with respective controls (Sen et al., 2018). 100 seconds of pre-scans and 1000 seconds of time-dependent enzyme kinetics were measured. At least 2 – 4 kinetic traces were collected per reaction. Data were normalized to their respective pre-scans to avoid problems that may be caused by variabilities between reactions. The time-dependent FRET signals were excited at 530 nm and measured at 670 nm. The data analysis was performed in the OriginLab software package.

#### ATPase activity measurements

7-Diethylamino-3-[N-(2-maleimidoethyl)-carbamoyl]-coumarin-conjugated phosphate binding protein A197C (MDCC-PBP) (Brune et al., 1994) is used to detect inorganic phosphate (P_i_) release from ATPase activity in real-time. Before the reaction, ATP was cleared of free P_i_ by performing a mopping reaction. In order to mop the ATP, 10 mM ATP was incubated with 1 U/mL PNPase (Sigma, N2415-100UN) and 200 uM 7-methylguanosine (Sigma, M0627-100MG) in mopping buffer (25 mM HEPES, 75 mM NaCl, 5 mM MgCl_2_, 1 mM DTT) for 2 hours at room temperature. ATPase assay reaction conditions were 2 nM SWI/SNF, 5 nM nucleosome, and 100 uM ATP in respective pH buffers; pH 6.5 (25 mM MES, 0.2 mM EDTA, 5 mM MgCl_2_, 70 mM KCl, 1 mM DTT), pH 7 (25 mM Tris, 0.2 mM EDTA, 5 mM MgCl_2_, 70 mM KCl, 1 mM DTT) or pH 7.6 (25 mM HEPES, 0.2 mM EDTA, 5 mM MgCl_2_, 70 mM KCl, 1 mM DTT). The measurements were performed on a Tecan Infinite 1000, with excitation at 405 nm and emission at 460 nm. Pre-scan measurements were taken to detect the basal level of signal per reaction. The time-dependent measurements were taken upon ATP addition, which started the reaction. At least 3-4 kinetic traces were analyzed using the steady-state equation using Graph Pad Prism 8 software.

### All-atom simulations

All-atom simulations were run with the ABSINTH implicit solvent model and CAMPARI Monte Carlo simulation (V2.0) (http://campari.sourceforge.net/) (85). The combination of ABSINTH and CAMPARI has been used to examine the conformational behavior of disordered proteins with good agreement to experiment (57, 86, 87).

All simulations were started from randomly generated non-overlapping random-coil conformations, with each independent simulations using a unique starting structure. Monte Carlo simulations perturb and evolve the system via a series of moves that alter backbone and sidechain dihedral angles, as well as rigid-body coordinates of both protein sequences and explicit ions. Simulation analysis was performed using CAMPARITraj (www.ctraj.com) and MDTraj (88).

ABSINTH simulations were performed with the ion parameters derived by Mao et al. and using the abs_opls_3.4.prm parameters (54). All simulations were run at 15 mM NaCl and 325 K, a simulation temperature previously shown to be a good proxy for *bona fide* ambient temperature (57, 89). A summary of the simulation input details is provided in **Supplemental Table 5**. For SNF5^71-120^ simulations twenty independent simulations were run for each combination of pH (as defined by histadine protonation state) and mutational state. For SNF5^195-223^, the high glutamine content made conformational sampling challenging, as has been observed in previous glutamine-rich systems, reflecting the tendancy for polyglutamine to undergo intramolecular chain collapse (90–92). To address this challenge we ran hundereds of short simulations (with a longer equilibration period than in SNF^71-120^) that are guaranteed to be uncorrlated due to their complete independence (93). Simulation code and details can be found at: https://github.com/holehouse-lab/supportingdata/tree/master/2021/Gutierrez_QLC_2021

### Bioinformatic analysis

All protein sequence analysis was performed with localCIDER, with FASTA files read by protfasta (https://github.com/holehouse-lab/protfasta) (94). Sequence alignments were performed using clustal omega (95). Sequence conservation was computed using default properties in with the score_conservation program as defined by Capra et al. (96). Proteomes were downloaded from UniProt (97).

Low-complexity sequences were identified using Wooton-Fedherhen complexity (98, 99). Sequence complexity is calculated over a sliding window size of 15 residues, and a threshold of 0.6 was used for binary classification of a residue as ‘low’ or ‘high’ complexity. After an initial sweep, gaps of up to 3 “high complexity residues” between regions of low-complexity residues were converted to low-complexity. Finally, contiguous stretches of 30 residues or longer were taken as the complete set of low-complexity regions in the proteome. The full set of those SEG-defined LCDs for human, drosophila, dictyostelium and cerevisiae proteomes is provided as FASTA files at: https://github.com/holehouse-lab/supportingdata/tree/master/2021/Gutierrez_QLC_2021/

## Acknowledgements

We thank Conor Howard for help with initial bioinformatics and conception of this project and Morgan Delarue for help with MatLab analysis. We thank David Truong, Sudarshan Pinglay and JoAnna Klein for help writing the manuscript; Ivan Tarride for help with figure design; and Karsten Weis, Jeremy Thorner, and Douglas Koshland for advice, strains, plasmids and reagents. We gratefully acknowledge funding from the William Bowes Fellows program, the Vilcek Foundation, and the HHMI HCIA summer institute (LJH); Becas Chile (JIG) and the National Science Foundation Graduate Research Fellows Program (GB).

## Author contributions

JIG and LJH designed the study. JIG carried out most experiments and wrote the initial paper. GB undertook RNA-seq analysis. YK and CLP undertook and analyzed *in vitro* SWI/SNF nucleosome remodeling experiments. ASH performed and analyzed all-atom Monte Carlo simulations and undertook sequence and evolutionary analyses. KT and AD purified SWI/SNF complexes. JIG and LJH wrote the final paper with contributions from GB, AD, ASH and CLP.

