## Supplemental Figures and Tables for "SWI/SNF senses carbon starvation with a pH-sensitive low complexity sequence"

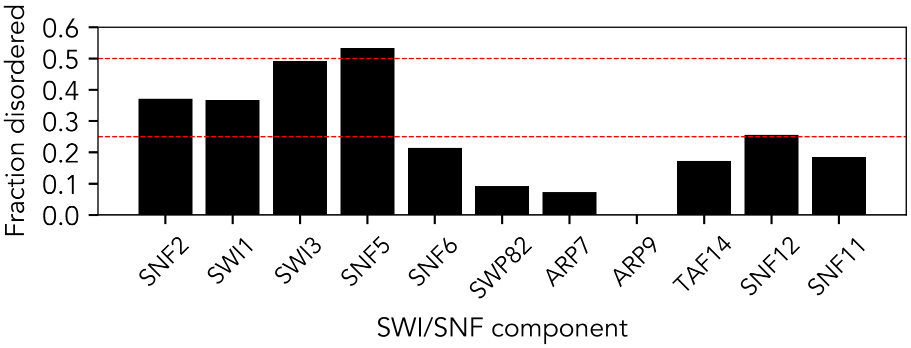


**Figure 1 – figure supplement 1: The SWI/SNF complex has 10/11 subunits with significant disorder.** Fractional disorder in each of the core eleven SWI/SNF components. Dashed red lines represent 25% and 50% disorder. Five of the eleven components contain over 25% disorder. Disorder prediction performed using MobiDBLite (see methods).

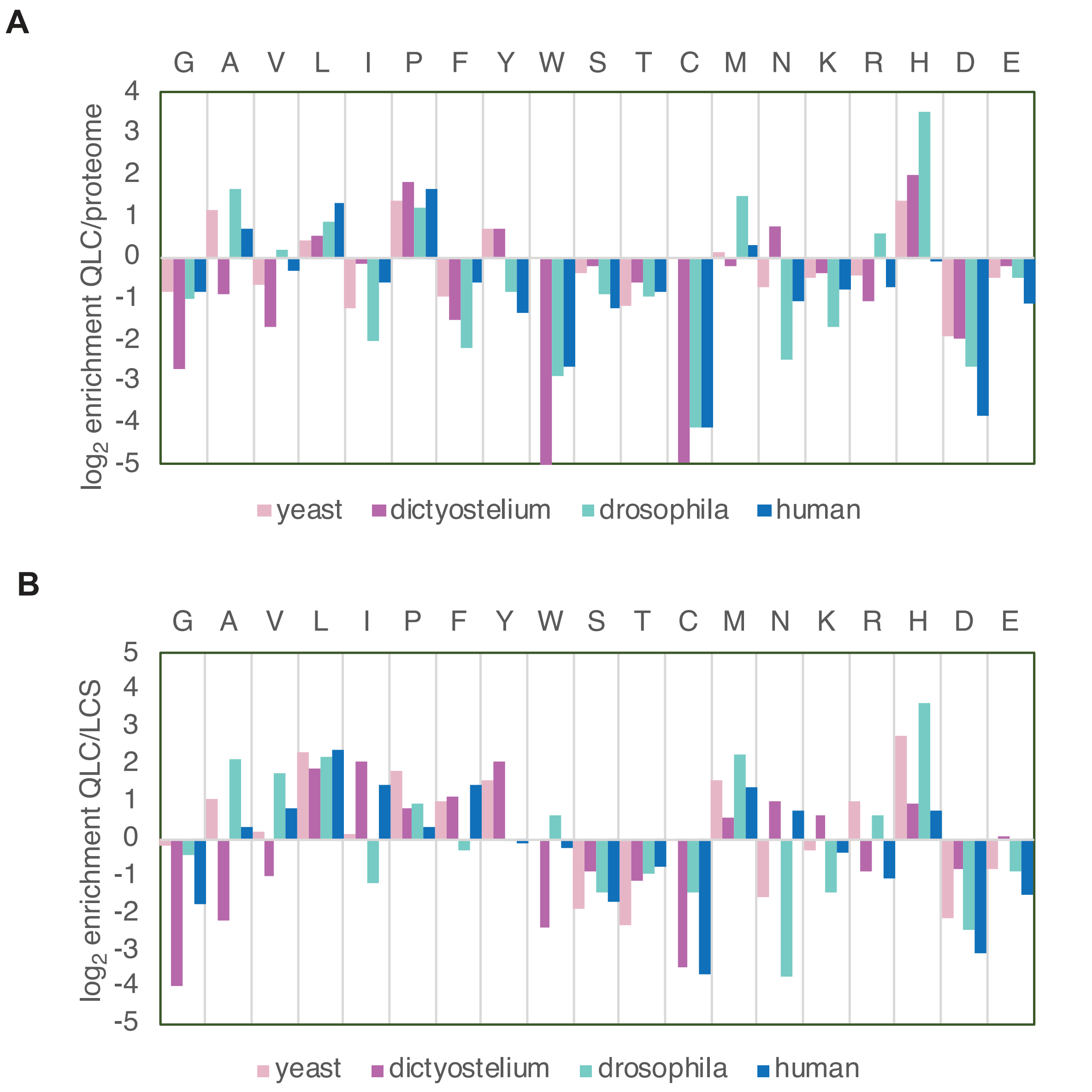


**Figure 1 – figure supplement 2: Histidines are enriched in glutamine-rich low-complexity sequences.** Amino-acid frequencies within glutamine-rich low-complexity sequences (QLCs) in *S. cerevisiae* (yeast), *Dictyostelium discoides*, *Drosophila melanogaster*, and humans. **A)** Enrichment of each amino acid in QLCs compared to global amino-acid frequencies in each proteome. **B)** Enrichment of each amino acid in QLCs compared to amino-acid frequencies in all low-complexity sequences identified using Wooton-Fedherhen complexity (see methods).

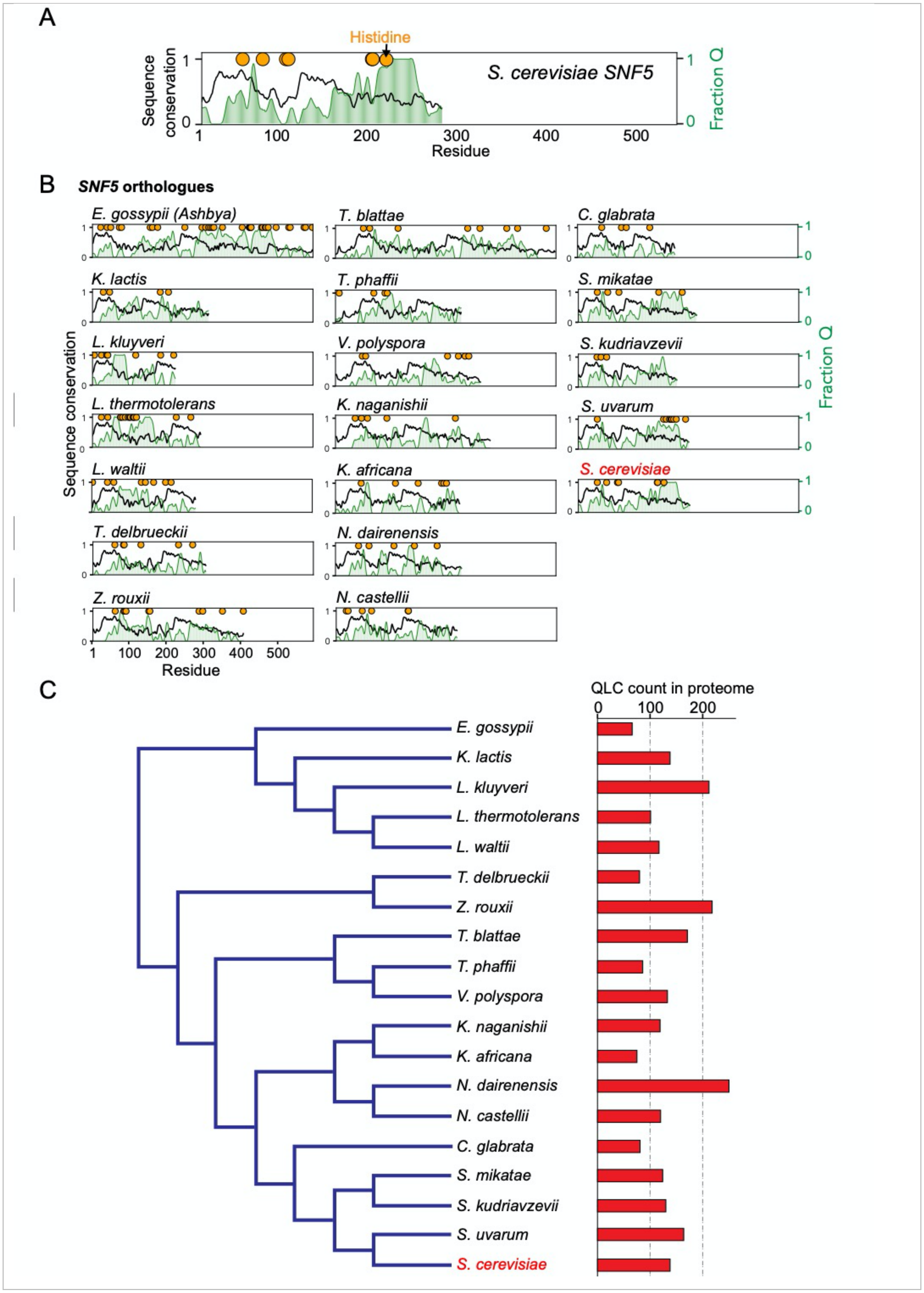

A
Histidine
S. cerevisiae SNF5
Sequence conservation
Fraction Q
Residue
B
SNF5 orthologues
E. gossypii (Ashbya)
T. blattae
C. glabrata
K. lactis
T. phaffii
S. mikatae
L. kluyveri
V. polyspora
S. kudriavzevii
L. thermotolerans
K. naganishii
S. uvarum
L. waltii
K. africana
S. cerevisiae
T. delbrueckii
N. dairenensis
Z. rouxii
N. castellii
C
QLC count in proteome

**Figure 1 – figure supplement 3: The *SNF5* N-terminal glutamine-rich low complexity domain (with embedded histidines) is broadly conserved across *ascomycota*. A)** Black line: Per-residue sequence conservation was assessed by Jensen-Shannon divergence of physicochemical properties across twenty distinct protein sequences (see methods). Conservation values of below 0.5 reflect poor conservation. There are two short islands of conservation between residues 19-54 and 122-161 of *S. cerevisiae SNF5*. Green area: Fraction of glutamine within the QLC. The seven histidine residues are shown in orange. **B)** The same analysis for 19 *ascomycetes*. **C)** Phylogeny of *ascomycetes* analyzed above, with the total number of QLCs identified in each proteome shown to the right.

**Supplemental Table 1: Comparison of sequence properties of *SNF5* N-terminal IDRs**

| Species | Number of His | Number of Gln | Total length |
|---|---|---|---|
| *V. polyspora* | 6 | 84 | 390 |
| *T. phaffii* | 4 | 107 | 338 |
| *T. blattae* | 8 | 187 | 590 |
| *N. dairenensis* | 5 | 92 | 339 |
| *N. castellii* | 6 | 77 | 325 |
| *K. naganishii* | 5 | 109 | 416 |
| *K. africana* | 6 | 88 | 334 |
| *C. glabrata* | 4 | 46 | 262 |
| *S. uvarum* | 9 | 119 | 297 |
| *S. kudriavzevii* | 3 | 89 | 265 |
| *S. mikatae* | 5 | 136 | 318 |
| *S. cerevisiae* | 7 | 125 | 300 |
| *Z. rouxii* | 10 | 122 | 405 |
| *T. delbrueckii* | 6 | 76 | 304 |
| *K. lactis* | 4 | 109 | 312 |
| *E. gossypii* | 34 | 253 | 593 |
| *E. cymbalariae* | 9 | 100 | 261 |
| *L. kluyveri* | 7 | 81 | 223 |
| *L. thermotolerans* | 15 | 119 | 291 |
| *L. waltii* | 8 | 91 | 279 |

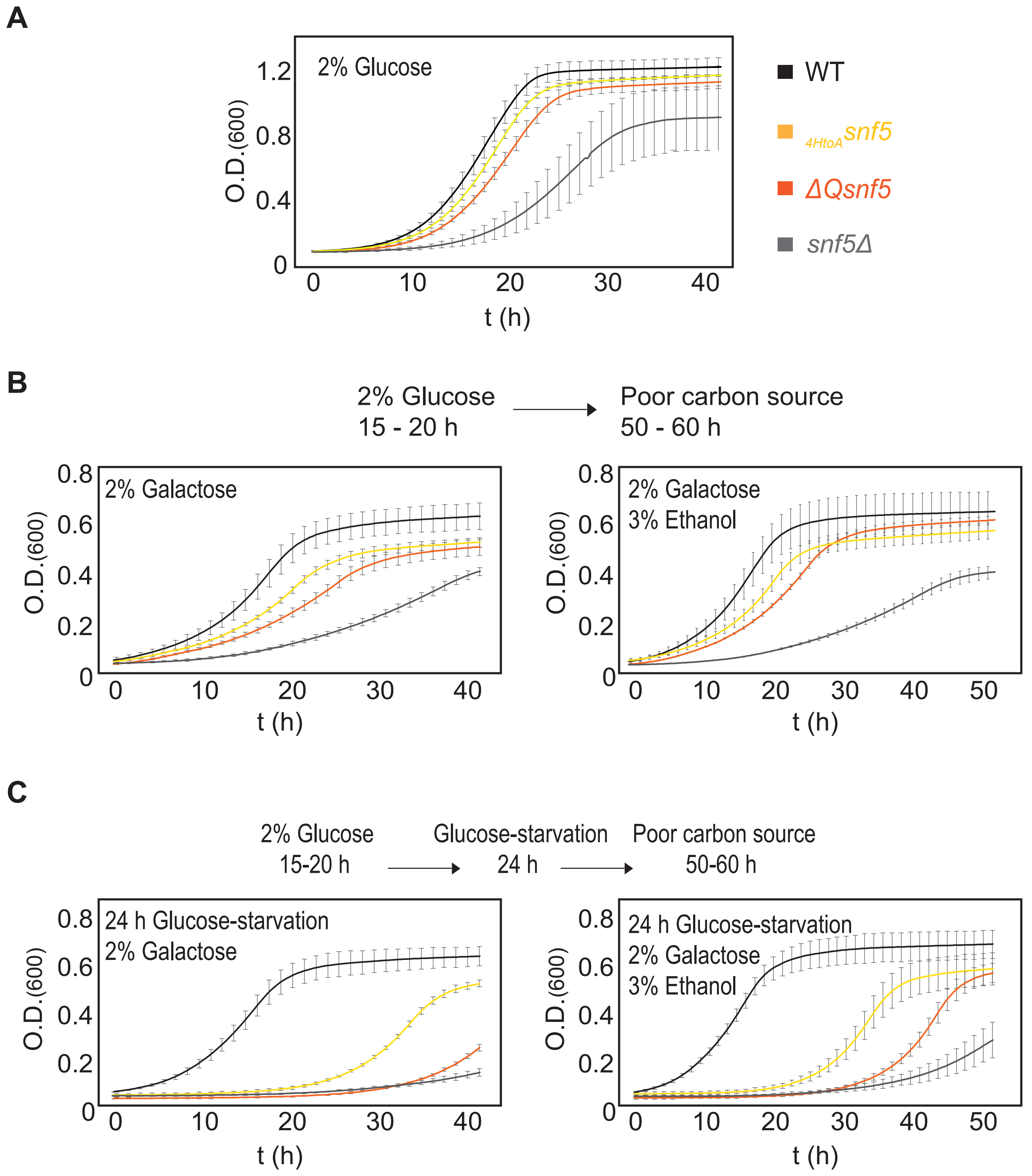


**Figure 1 – figure supplement 4: The *SNF5* QLC is important for recovery from carbon starvation.** Growth rate was assessed in a plate reader in various conditions. **A)** Comparison of growth rate of WT, *ΔQsnf5*, $_{HtoA}$*snf5*, and *snf5Δ* strains in synthetic complete media with glucose. **B)** Cells were immediately switched from glucose to poor carbon sources. **C)** Cells were subjected to acute starvation for 24 h and then switched to poor carbon sources.

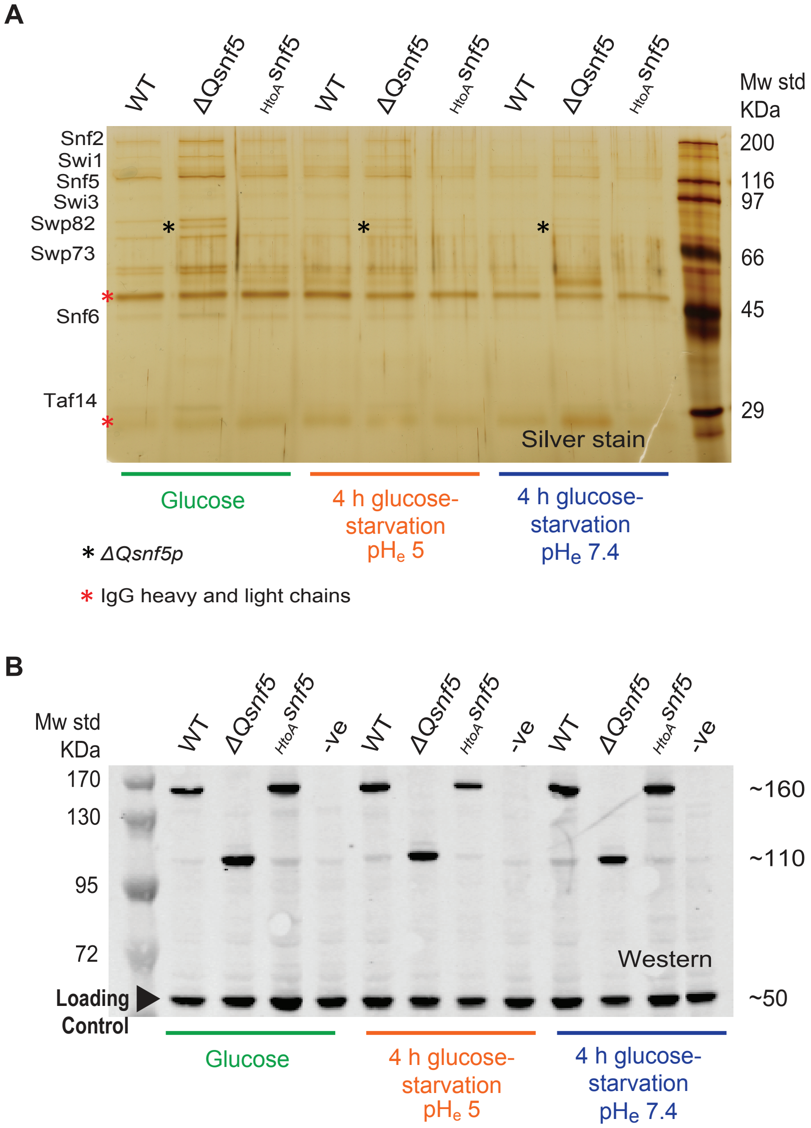


**Figure 1 – figure supplement 5: Mutation of the *SNF5* QLC does not lead to protein degradation or loss of SWI/SNF complex integrity. A)** *The entire SWI/SNF complex copurifies with SNF2 in all strains and conditions*. The endogenous *SNF2* gene was TAP-tagged, and used to immunoprecipitate the SWI/SNF complex from WT, *ΔQsnf5*, or $_{HtoA}$*snf5* strains either exponentially growing in glucose, or after 4 h acute carbon starvation in media titrated to $pH_e$ 5 or 7.4 (indicated at bottom). A silver stain of an SDS-PAGE analysis is shown. **B)** *Neither SNF5 nor its mutant alleles are degraded upon glucose-starvation.* Western blots of the TAP-tagged *SNF5* alleles in various conditions (indicated at bottom). TAP-tagged *ΔQ-snf5*

runs at ~110 KDa, 288 amino acids smaller than WT (~160 KDa). An anti-glucokinase antibody was used as a loading control (Bottom band at ~50 kDa).

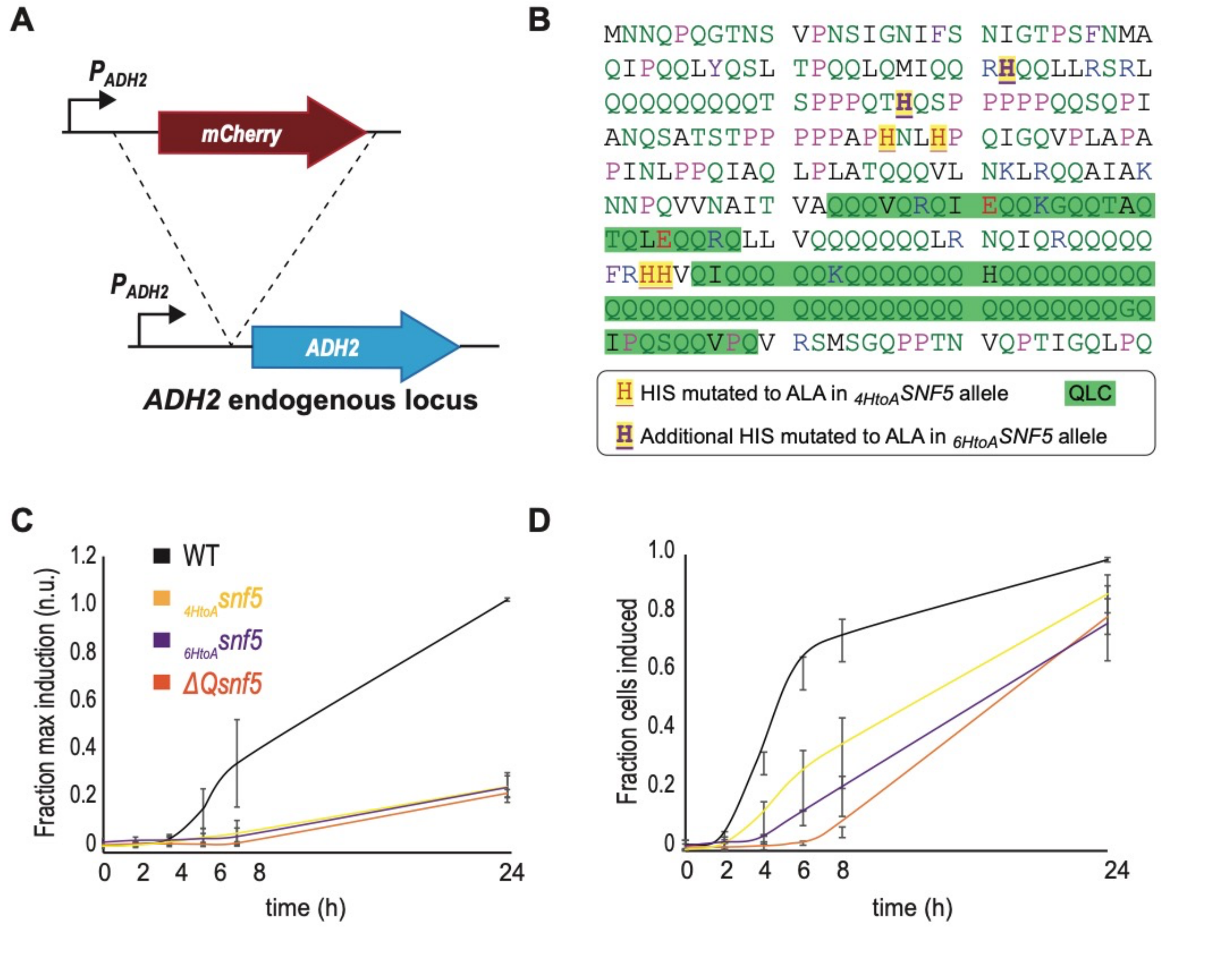


**Figure 1 – figure supplement 6: The *SNF5* QLC and embedded histidines are required for efficient *ADH2* induction upon carbon starvation. A)** Schematic of the $P_{ADH2}$-*mCherry* reporter gene: the reporter construct was integrated into the endogenous *ADH2* locus resulting in a tandem repeat of the reporter gene followed and an intact *ADH2* gene. **B)** Sequence of the *SNF5* N-terminus with the 4/7 histidines that were mutated in the $_{4HtoA}$*snf5* allele highlighted as red on yellow, and the additional 2 histidines that were mutated in the $_{6HtoA}$*snf5* allele highlighted as purple on yellow. **C)** $P_{ADH2}$-*mCherry* induction during carbon starvation assessed by fluorescence cytometry, normalized to the maximal induction (median mCherry fluorescence at 24 h in *SNF5* WT strains). **D)** The fraction of the cells that induce $P_{ADH2}$-*mCherry* induction at each time point during carbon starvation (see methods). The *SNF5* alleles compared are: WT, *ΔQsnf5*, $_{4HtoA}$*snf5* (referred to in the rest of the manuscript as simply $_{HtoA}$*snf5*) and the $_{6HtoA}$*snf5* strains with an additional 2 histidines (6/7 total) mutated to alanine. There is no significant difference between the ${}_{4HtoA}$*snf5* and ${}_{6HtoA}$*snf5* strains in these experiments. Mean and standard deviation are shown in each plot.

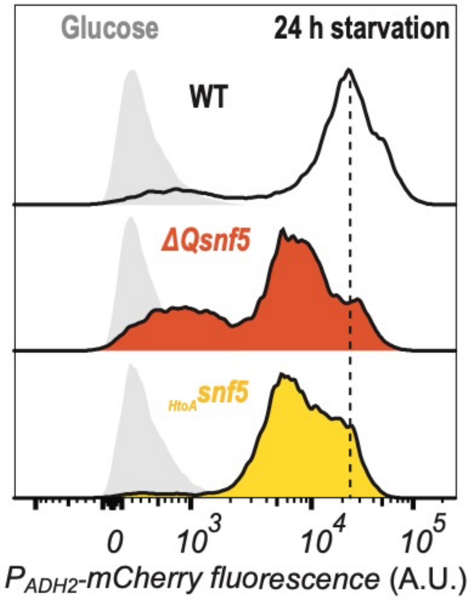


**Figure 2 – figure supplement 1: All strains ultimately express some amount of *ADH2*.** Cytometry data showing $P_{ADH2}$-*mCherry* induction either in glucose (light grey peaks to left) or after 24 h of acute carbon starvation (dark lines, and color-coded by strain).

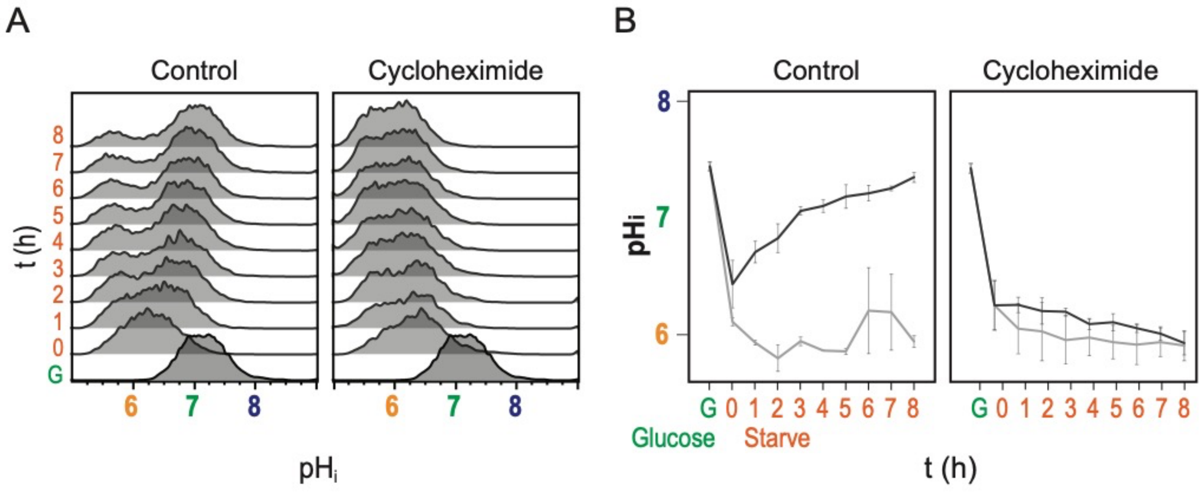


**Figure 2 – figure supplement 2: Recovery of $pH_i$ requires new protein translation.**

**A)** Cytometry data showing nucleocytoplasmic pH ($pH_i$), calculated using the ratiometric pHluorin probe. **B)** Quantification of $pH_i$ data (see methods), two populations are detected and indicated in the black and grey lines. Mean and standard deviation of three biological replicates are plotted. Cells were switched to acute carbon starvation media titrated to the optimal $pH_e$ of 5.5 at time 0, but the right panels show cells additionally exposed to the translational inhibitor cycloheximide. Intracellular pH fails to recover without new protein translation.

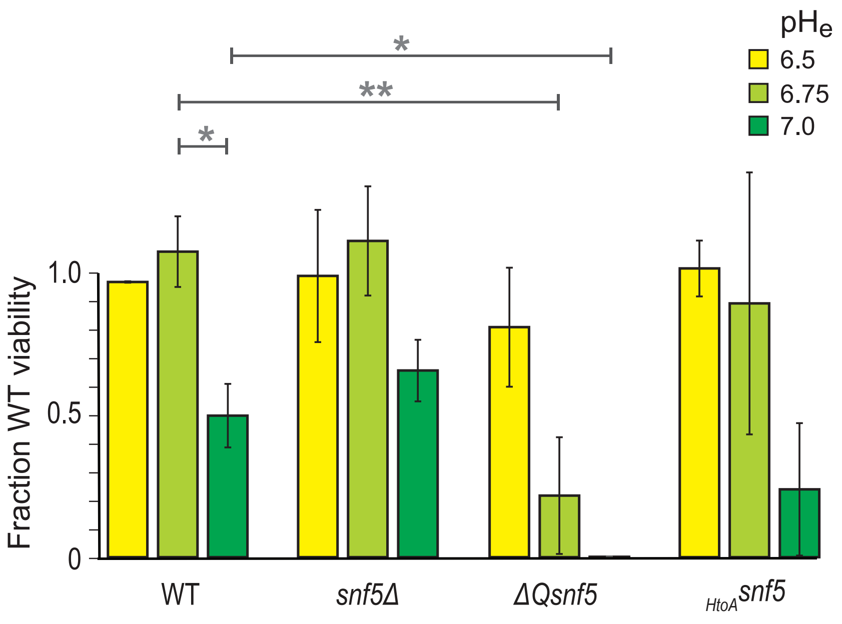


**Figure 3 – figure supplement 1: Deletion of the N-terminal glutamine rich domain of *SNF5* renders cells hypersensitive to starvation at suboptimal extracellular pH.** Cells were grown to log phase and then subjected to acute carbon starvation in media titrated to various $pH_e$ values (see legend). After 24 h starvation, cells were plated to determine the number of colony-forming units compared to WT cells starved at $pH_e$ 6.5. Mean and standard deviation of 3 biological replicates are shown. Single and double asterisks represent $p < 0.05$ and $p < 0.01$ respectively from t-tests.

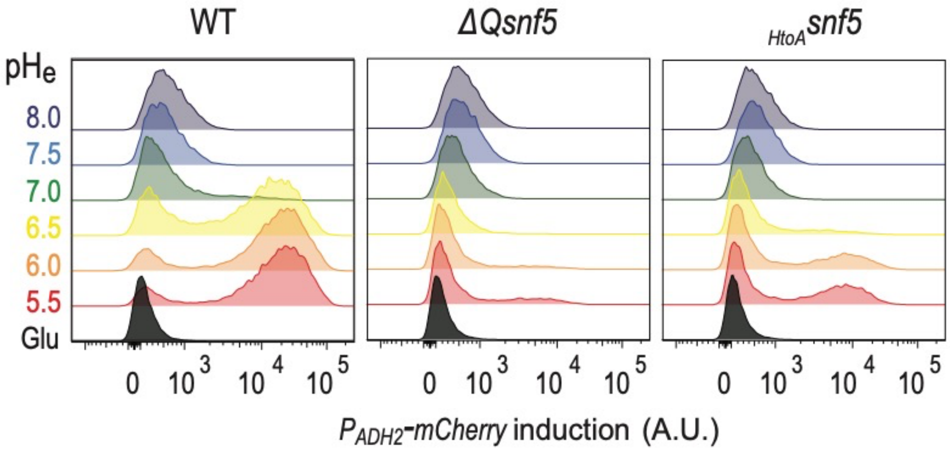


**Figure 3 – figure supplement 2: *$P_{ADH2}$-mCherry* induction requires an acidic extracellular environment and the *SNF5* QLC.** Cytometry data showing expression levels of the *$P_{ADH2}$-mCherry* reporter from WT, *ΔQsnf5*, or *$_{HtoA}$snf5* cells either growing in glucose (Glu), or 6 h after acute carbon starvation in media titrated to various $pH_e$ values (these are representative source data for Figure 3A).

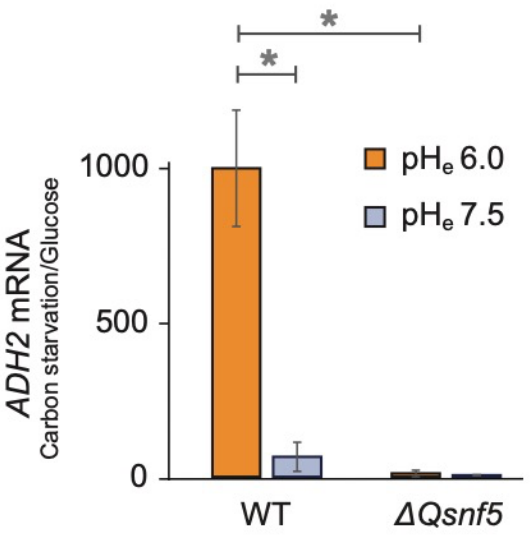


**Figure 3 – figure supplement 3: Expression of the endogenous *ADH2* mRNA requires an acidic extracellular environment and the *SNF5* QLC.** RT-qPCR data showing *ADH2* mRNA levels. The ratio of *ADH2* levels in carbon-starved cells to cells growing in glucose is shown. *ACT1* was used as an internal control to normalize *ADH2* values. WT and *ΔQsnf5* strains were carbon starved in media titrated to $pH_e$ of either 6.0 or 7.5. Mean and standard deviation of three biological replicates are shown. Single and double asterisks represent $p < 0.05$ and $p < 0.01$ respectively from t-tests.

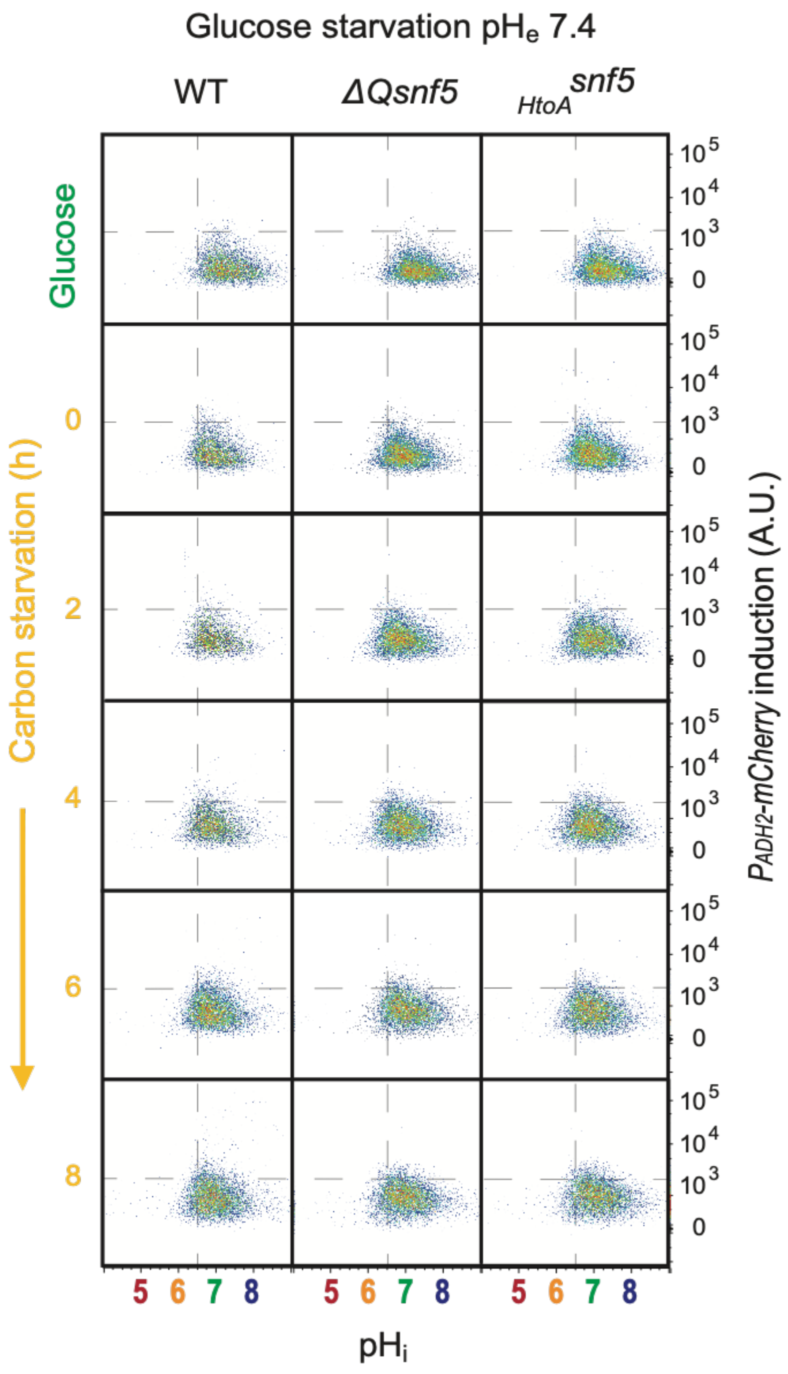


**Figure 3 – figure supplement 4: Transient acidification of cells requires an acidic extracellular environment.** Flow cytometry for WT, *ΔQsnf5*, or $_{HtoA}$*snf5* strains: the **x-axis** shows nucleocytosoplasmic pH ($pH_i$), while the **y-axis** shows fluorescence from the $P_{ADH2}$-*mCherry* reporter. Panels show cells grown in glucose (top) and then (2nd to bottom) after 0 - 8 h of acute glucose-starvation.

| Group | TF | Fraction group promoters | Fraction promoters in total genome | p value | Target promoters |
|---|---|---|---|---|---|
| | Gis1 | 0.67 | 0.05 | < 1E-13 | CIT2, ADY2, NDE2, CTA1, STL1, ICL1, HSP12, SPG1, PEX18, POT1, RGI2, JEN1, PCK1, IDP2, FBP1, CAT2, SPG4, SIP18, ADH2, ALP1, MLS1, LPX1, CRC1, GRE1, CIT3, PDH1, YAT1, ECM13 |
| | Adr1 | 0.64 | 0.03 | < 1E-13 | ADY2, CTA1, ATO3, STL1, HSP12, POX1, SPG1, ARO9, POT1, RGI2, JEN1, FOX2, IDP2, FBP1, CAT2, SPG4, ADH2, ALP1, SPS19, MLS1, LPX1, CRC1, CIT3, PDH1, ACS1, ECM13, ARG3 |
| | Cat8 | 0.60 | 0.14 | < 1E-13 | CIT2, ADY2, CTA1, ATO3, STL1, YAT2, ICL1, POX1, SPG1, ARO9, POT1, SFC1, JEN1, FOX2, PCK1, IDP2, FBP1, CAT2, ADH2, MLS1, LPX1, CRC1, ACS1, YAT1, SIP4 |
| | Bas1 | 0.93 | 0.01 | 2E-13 | CIT2, ADY2, NDE2, CTA1, ATO3, STL1, YAT2, ICL1, HSP12, POX1, SPG1, ARO9, PEX18, POT1, RGI2, SFC1, JEN1, PCK1, IDP2, FBP1, CAT2, SPG4, SIP18, ADH2, ALP1, SPS19, MLS1, LPX1, CRC1, GRE1, CIT3, PDH1, ACS1, YAT1, ECM13, SIP4, ARG3, DPA10, YAR035C-A |
| | Oaf1 | 0.55 | 0.03 | 1E-11 | CIT2, ADY2, CTA1, ATO3, STL1, YAT2, ICL1, POX1, PEX18, POT1, FOX2, PCK1, SPG4, SIP18, ADH2, ALP1, SPS19, MLS1, LPX1, CRC1, GRE1, CIT3, YAT1 |
| | Yox1 | 0.55 | 0.02 | 3E-10 | CIT2, ADY2, NDE2, CTA1, STL1, YAT2, ICL1, HSP12, SPG1, PEX18, POT1, RGI2, JEN1, PCK1, CAT2, SPG4, ADH2, MLS1, GRE1, ACS1, YAT1, SIP4, DPA10 |
| | Xbp1 | 0.64 | 0.02 | 4E-10 | CIT2, ADY2, NDE2, CTA1, ATO3, STL1, ICL1, HSP12, POX1, ARO9, PEX18, POT1, RGI2, SFC1, FOX2, PCK1, IDP2, FBP1, CAT2, SPG4, MLS1, CRC1, CIT3, PDH1, YAT1, DPA10, YAR035C-A |
| | Gln3 | 0.79 | 0.01 | 4E-10 | CIT2, NDE2, CTA1, ATO3, STL1, YAT2, ICL1, HSP12, SPG1, ARO9, PEX18, POT1, RGI2, JEN1, PCK1, IDP2, FBP1, CAT2, SPG4, SIP18, ADH2, SPS19, MLS1, LPX1, CRC1, IRC13, CIT3, PDH1, ACS1, YAT1, ECM13, SIP4, ARG3 |
| | Sok2 | 0.74 | 0.01 | 5E-09 | CIT2, ADY2, ATO3, STL1, YAT2, ICL1, HSP12, POX1, SPG1, PEX18, POT1, RGI2, SFC1, JEN1, FOX2, FBP1, CAT2, SPG4, SIP18, ADH2, SPS19, MLS1, CRC1, GRE1, CIT3, PDH1, ACS1, YAT1, ECM13, SIP4, ARG3 |
| | Rme1 | 0.38 | 0.02 | 5E-08 | ADY2, ICL1, HSP12, POX1, ARO9, PEX18, SFC1, JEN1, FOX2, FBP1, CAT2, SPG4, GRE1, CIT3, ACS1, SIP4 |
| 1 | Hac1 | 0.55 | 0.02 | 2E-07 | ADY2, NDE2, CTA1, ATO3, STL1, YAT2, ICL1, HSP12, SPG1, PEX18, POT1, RGI2, JEN1, FOX2, IDP2, ADH2, SPS19, MLS1, CRC1, IRC13, YAT1, SIP4, ARG3 |
| | Sfl1 | 0.33 | 0.02 | 1E-06 | ADY2, NDE2, CTA1, ATO3, STL1, YAT2, ICL1, HSP12, SPG1, PEX18, POT1, RGI2, JEN1, FOX2, IDP2, ADH2, SPS19, MLS1, CRC1, IRC13, YAT1, SIP4, ARG3 |
| | Hot1 | 0.14 | 0.06 | 2E-06 | STL1, ICL1, HSP12, SPG4, SIP18, GRE1 |
| | Gat4 | 0.26 | 0.03 | 2E-06 | NDE2, HSP12, ARO9, PEX18, SFC1, JEN1, PCK1, CAT2, SPG4, LPX1, ECM13 |
| | Fhl1 | 0.60 | 0.01 | 3E-06 | CIT2, ADY2, NDE2, STL1, HSP12, POX1, SPG1, ARO9, PEX18, POT1, RGI2, SFC1, JEN1, FOX2, PCK1, SPG4, SIP18, SPS19, MLS1, CRC1, GRE1, CIT3, ACS1, SIP4, ARG3 |
| | Tog1 | 0.19 | 0.04 | 4E-06 | ICL1, POX1, POT1, FOX2, PCK1, IDP2, FBP1, MLS1 |
| | Hsf1 | 0.55 | 0.01 | 1E-05 | CIT2, ADY2, NDE2, ATO3, STL1, HSP12, POX1, ARO9, RGI2, SFC1, IDP2, CAT2, SPG4, SIP18, ADH2, SPS19, MLS1, CRC1, PDH1, ACS1, ECM13, SIP4, ARG3 |
| | Ixr1 | 0.64 | 0.01 | 2E-05 | ADY2, NDE2, CTA1, ATO3, STL1, YAT2, HSP12, SPG1, ARO9, RGI2, SFC1, FOX2, IDP2, FBP1, CAT2, SIP18, ADH2, ALP1, MLS1, YNL013C, CRC1, GRE1, ACS1, YAT1, ECM13, SIP4, YAR035C-A |
| | Rgm1 | 0.29 | 0.02 | 2E-05 | STL1, HSP12, POX1, ARO9, SFC1, PCK1, IDP2, FBP1, CAT2, LPX1, CIT3, PDH1 |
| | Ume6 | 0.48 | 0.01 | 2E-05 | CIT2, ADY2, STL1, HSP12, SPG1, ARO9, SFC1, FOX2, FBP1, CAT2, SPG4, SIP18, ALP1, CRC1, PDH1, ACS1, YAT1, ECM13, SIP4, ARG3 |
| | Msn4 | 0.64 | 0.01 | 2E-05 | CIT2, NDE2, CTA1, STL1, YAT2, ICL1, HSP12, POX1, ARO9, PEX18, POT1, RGI2, FOX2, PCK1, IDP2, CAT2, SPG4, SIP18, SPS19, MLS1, LPX1, CRC1, GRE1, CIT3, PDH1, YAT1, SIP4 |
| | Sip4 | 0.17 | 0.03 | 4E-05 | SFC1, PCK1, FBP1, ADH2, MLS1, ACS1, SIP4 |
| | Mig3 | 0.50 | 0.01 | 4E-05 | CIT2, NDE2, CTA1, ATO3, STL1, HSP12, POX1, ARO9, PEX18, POT1, SFC1, JEN1, PCK1, FBP1, CAT2, SPG4, SIP18, CRC1, IRC13, ACS1, YAT1 |
| | Gat4 | 0.32 | 0.05 | 3E-11 | HSP30, YCR102C, TRR1, TSA2, QCR7, COX13, TRX2, OM45, REE1, YLR012C, YMR090W, PAI3, HOR7, YNL134C, TMA16, OYE3, SSA3, HSP26, TDH1 |
| | Pho2 | 0.44 | 0.02 | 1E-08 | CHA1, MIC10, YCR102C, YDR149C, TRR1, TSA2, QCR7, COX13, CGR1, STF2, NQM1, CUP1-1, CUP1-2, SPL2, OM45, BAT2, SRX1, YKL030W, YLR460C, PAI3, HOR7, YNL134C, EGO4, YOR338W, TDH1, LYS20 |
| | Mig3 | 0.51 | 0.02 | 1E-06 | HSP30, YCR102C, TRR1, TSA2, CGR1, NQM1, TRX2, CUP1-1, CUP1-2, SPL2, YIL168W, SDL1, BAT2, SRX1, CIS1, YLR460C, YMR090W, PAI3, SUI1, YNL194C, YNL134C, EGO4, TMA16, YOR338W, EEB1, AI1, AI2, HSP26, LYS20, YMR046C |
| 2 | Tbs1 | 0.12 | 0.07 | 2E-06 | YCR102C, SPL2, REE1, YLR460C, ATP18, YNL134C, LYS20 |
| | Gis1 | 0.27 | 0.03 | 3E-06 | TSA2, QCR7, COX13, STF2, NQM1, TRX2, OM45, REE1, YLR012C, YMR090W, PAI3, YNL194C, YNL134C, SSA3, HSP26, TDH1 |
| | Hsf1 | 0.49 | 0.02 | 2E-05 | CHA1, GRX1, HSP30, YCR102C, TSA2, COX13, STF2, NQM1, CUP1-1, CUP1-2, SPL2, REE1, BAT2, SRX1, YKL030W, YMR090W, PAI3, HOR7, SUI1, YNL194C, YNL134C, YOR338W, OYE3, SSA3, HSP26, TDH1, LYS20, YBL005W-A, YMR046C |
| | Fhl1 | 0.51 | 0.02 | 3E-05 | CHA1, GRX1, HSP30, TSA2, QCR7, COX13, STF2, NQM1, CUP1-1, CUP1-2, SPL2, YIL168W, OM45, REE1, YLR012C, YMR090W, PAI3, HOR7, SUI1, YNL194C, YNL134C, EGO4, RRS1, YOR338W, OYE3, AI1, HSP26, TDH1, COQ21, LYS20 |
| | Arr1 | 0.56 | 0.03 | 3E-12 | VID24, FUS1, MET32, HXT7, DSF1, GSY1, MIG2, RCK1, MUP1, IMA1, YHR022C, HXT4, OPT1, YJR115W, MSN4, PHD1, UGP1, YKR075C, MHT1, MMP1, ICT1, TIS11, HXT2, MET2, YNR014W, MAN2, SAM3, BNA3 |
| | Mig2 | 0.26 | 0.04 | 1E-08 | EMI2, GSY1, FMP48, IMA1, HXT4, YHR210C, SUC2, RPI1, OPT1, YJR115W, ICT1, HXT2, MAN2 |
| | Yh1 | 0.44 | 0.02 | 3E-08 | VID24, FUS1, HXT7, EMI2, GSY1, MIG2, RCK1, FMP48, IMA1, OPT1, MSN4, PHD1, MMP1, HSP104, SSA2, ICT1, HXT2, FET3, YNR014W, MAN2, PFK27, CLN3 |
| | Opi1 | 0.26 | 0.03 | 2E-07 | FUS1, HXT7, LPP1, EMI2, MIG2, HXT4, MET3, PHD1, UGP1, MMP1, ICT1, TIS11, SAM3 |
| | Gis1 | 0.32 | 0.03 | 2E-07 | HXT7, RCK1, FMP48, MUP1, SPR3, IMA1, YHR022C, YHR210C, SUC2, MSN4, YKR075C, MMP1, TIS11, HXT2, YNR014W, MAN2 |
| | Rox1 | 0.40 | 0.02 | 2E-07 | VID24, FUS1, HXT7, EMI2, GSY1, MIG2, RCK1, IMA1, YHR210C, SUC2, RPI1, PHD1, UGP1, SSA2, ICT1, TIS11, HXT2, FET3, YMR320W, YNR014W |
| | Sfl1 | 0.32 | 0.03 | 6E-07 | FUS1, DSF1, RCK1, FMP48, OPT1, YJR115W, MSN4, PHD1, YKR075C, HSP104, ICT1, TIS11, FET3, YNR068C, MAN2, BNA3 |
| | Hac1 | 0.48 | 0.02 | 3E-06 | VID24, FUS1, HXT7, LPP1, EMI2, DSF1, GSY1, STR3, RCK1, YHR022C, HXT4, YHR210C, RPI1, MET28, MET3, YJR115W, ICT1, FET3, YMR320W, MET2, YNR014W, YNR068C, MAN2, BNA3 |
| | Yrr1 | 0.56 | 0.01 | 3E-06 | HXT7, EMI2, GSY1, MIG2, STR3, MUP1, HXT4, OPT1, MET3, MSN4, PHD1, UGP1, YKR075C, MHT1, MMP1, HSP104, SSA2, ICT1, TIS11, HXT2, FET3, MET2, YNR014W, YNR068C, MAN2, PFK27, SAM3, BNA3 |
| | Yox1 | 0.40 | 0.02 | 4E-06 | VID24, FUS1, MET32, GSY1, MIG2, RCK1, FMP48, IMA1, YHR210C, SUC2, OPT1, MSN4, PHD1, MMP1, SSA2, ICT1, HXT2, YNR014W, MAN2, CLN3 |
| 3 | Aft1 | 0.42 | 0.02 | 4E-06 | VID24, MET32, HXT7, EMI2, GSY1, SPR3, IMA1, HXT4, MET28, MET3, YJR115W, MSN4, UGP1, YKR075C, HSP104, ICT1, TIS11, HXT2, FET3, YMR320W, YNR014W |
| | Mig1 | 0.28 | 0.02 | 6E-06 | FUS1, HXT7, GSY1, FMP48, IMA1, YHR022C, HXT4, SUC2, RPI1, PHD1, MMP1, ICT1, HXT2, YNR014W |
| | Mig3 | 0.50 | 0.02 | 1E-05 | FUS1, MET32, HXT7, GSY1, MIG2, STR3, FMP48, IMA1, MET28, OPT1, MSN4, PHD1, YKR075C, MHT1, MMP1, HSP104, SSA2, ICT1, TIS11, HXT2, FET3, MET2, YNR068C, MAN2, PFK27 |
| | Sok2 | 0.60 | 0.01 | 1E-05 | VID24, FUS1, YDR249C, HXT7, EMI2, GSY1, MIG2, RCK1, FMP48, SPR3, YHR022C, HXT4, SUC2, RPI1, MET28, OPT1, YJR115W, MSN4, PHD1, HSP104, SSA2, ICT1, HXT2, FET3, MET2, YNR068C, MAN2, PFK27, IRC10, SAM3 |
| | Gzf3 | 0.24 | 0.03 | 1E-05 | VID24, FUS1, FMP48, SUC2, YKR075C, MMP1, SSA2, ICT1, TIS11, YNR014W, MAN2, BNA3 |
| | Ixr1 | 0.62 | 0.01 | 1E-05 | MET32, HXT7, PLM2, DSF1, GSY1, STR3, RCK1, FMP48, MUP1, YHR022C, HXT4, YHR210C, SUC2, RPI1, MET28, OPT1, MET3, MSN4, UGP1, YKR075C, MHT1, SSA2, HXT2, MET2, YNR068C, MAN2, YOL134C, IRC10, SAM3, CLN3, BNA3 |
| | Mot3 | 0.34 | 0.02 | 2E-05 | FUS1, HXT7, GSY1, FMP48, IMA1, YHR022C, HXT4, SUC2, RPI1, PHD1, HSP104, HXT2, FET3, MAN2, PFK27, SAM3, YPR050C |
| | Mga1 | 0.34 | 0.02 | 2E-05 | FUS1, HXT7, DSF1, YHR022C, HXT4, YHR210C, OPT1, MSN4, YKR075C, MHT1, HSP104, SSA2, ICT1, TIS11, MET2, YNR014W, MAN2 |
| | Rme1 | 0.28 | 0.02 | 2E-05 | VID24, FUS1, MET32, STR3, IMA1, SUC2, RPI1, MET3, PHD1, FET3, MET2, YNR014W, PFK27, SAM3 |
| | Cup2 | 0.30 | 0.02 | 3E-05 | HXT7, DSF1, GSY1, FMP48, MUP1, YHR022C, HXT4, RPI1, HSP104, ICT1, TIS11, HXT2, FET3, YNR068C, CLN3 |
| | Rgm1 | 0.26 | 0.02 | 3E-05 | HXT7, GSY1, HXT4, SUC2, RPI1, MSN4, HSP104, ICT1, FET3, YNR068C, MAN2, PFK27, IRC10 |
| | Stp1 | 0.32 | 0.02 | 5E-05 | VID24, EMI2, STR3, MUP1, RPI1, MET28, OPT1, MET3, PHD1, UGP1, MMP1, ICT1, TIS11, YMR320W, MET2, IRC10 |
| 4 | Cst6 | 0.88 | 0.00 | 2E-05 | ARO4, HXT3, HXK2, SCW4, HXT1, YJL218W, GPM1, KTI12, SHM2, RPS31, RPP0, ADE17, MF(ALPHA)1, YBL111C |
| | Rgt1 | 0.38 | 0.01 | 3E-05 | HXT3, HXK2, SCW4, HXT1, RPP0, MF(ALPHA)1 |

**Supplemental table 2: Transcription factors enriched in each gene group from RNA-seq analysis.** The YEASTRACT server used to find transcription factors enriched within the promoters of each of four gene sets defined by hierarchical clustering of genes significantly regulated upon carbon starvation (see figure 4E). YEASTRACT search settings were: DNA binding plus expression evidence; TF acting as either activator or inhibitor.

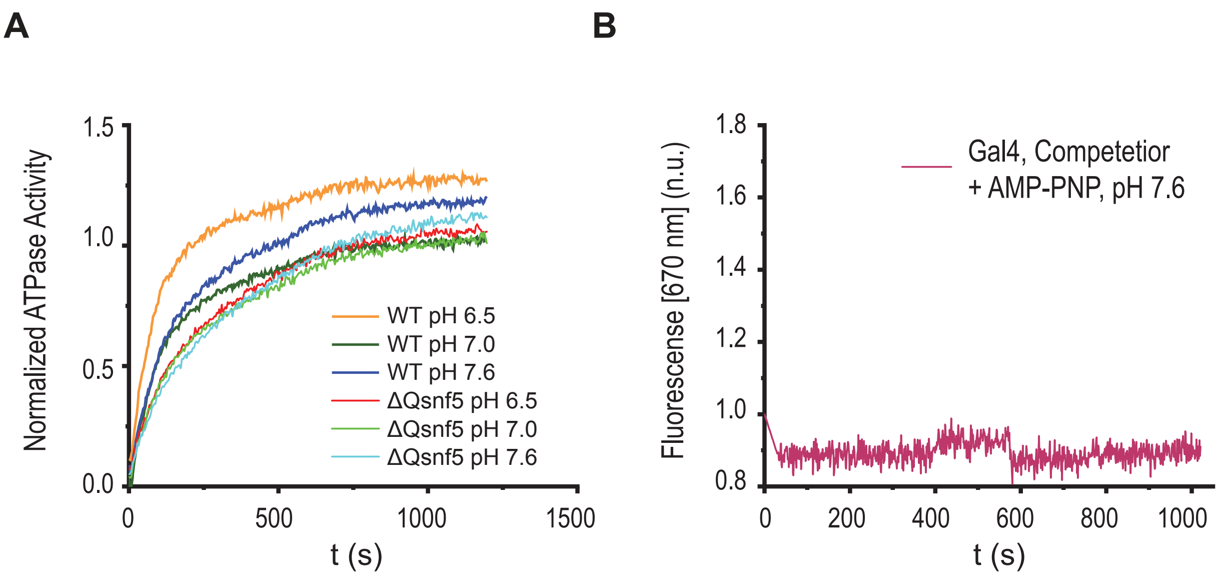


**Figure 5 – figure supplement 1: Basal ATPase activity is not affected by pH, and FRET changes require ATP hydrolysis. A)** ATPase activity of WT and *ΔQsnf5* mutant SWI/SNF complex in response to varied environmental pH. WT and mutant complexes do not show significant changes in ATPase activity (as assessed by inorganic phosphate release, see methods). **(B)** Representative kinetic trace for wildtype type SWI/SNF under recruitment conditions after addition of AMP-PNP (a non-hydrolysable ATP analogue).

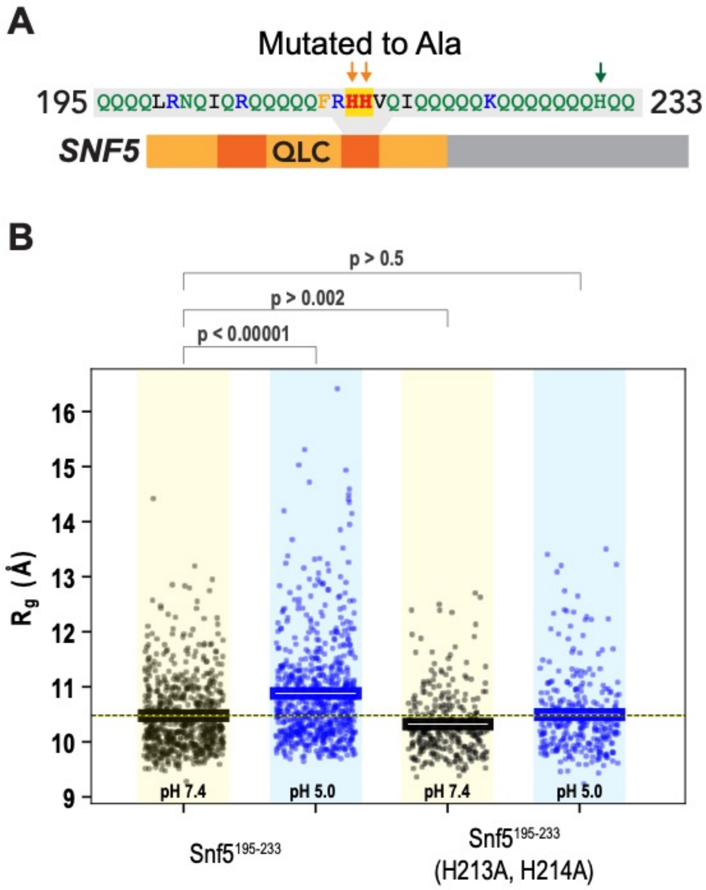


**Figure 6 – figure supplement 1: A second peptide within the N-terminal QLC of *SNF5* undergoes conformational expansion upon protonation. A)** Schematic of the *SNF5* gene, with the sequence and location of the simulated peptide indicated. **B)** Radius of gyration ($R_g$, **y-axis**) of all-atom Monte-Carlo simulations of amino acids 195-233 of the *SNF5* QLC with histidines either neutral (pH 7.4) or protonated (pH 5.0). Left two datasets are for the native peptide, right two datasets are with 2/3 histidines (H213 and H214) replaced with alanine, mimicking the $_{HtoA}$*SNF5* allele. Points represent the mean $R_g$ from all conformations sampled in each independent simulation (beginning from distinct random initial conformers). Bars represent the mean values of all simulations. P-values are from two-sided independent t-tests.

### Supplemental Table 4: SNF5 subregions examined by all-atom Monte Carlo simulations

| **Gene** | **Start** | **End** | **Sequence** |
|---|---|---|---|
| SNF5 (WT) | 71 | 120 | SPPPQTHQSPPPPPQQSQPIANQSATSTPPPPPAP**H**NL**H**PQIGQVPLAPA |
| SNF5 (H2A) | 71 | 120 | SPPPQTHQSPPPPPQQSQPIANQSATSTPPPPPAP**A**NL**A**PQIGQVPLAPA |
| SNF5 (WT) | 195 | 233 | QQQQLRNQIQRQQQQQFR**HH**VQIQQQQQKQQQQQQQHQQ |
| SNF5 (H2A) | 195 | 233 | QQQQLRNQIQRQQQQQFR**AA**VQIQQQQQKQQQQQQQHQQ |

**Supplemental Table 5:** Parameters used for all-atom Monte Carlo simulations

| Name | Droplet radius (Å) | Number of replicas | Steps per replica | Equilibration steps | Total ensemble |
|---|---|---|---|---|---|
| $SNF5^{71-120}$ WT | 94 | 20 | 125,000,000 | 5,000,000 | 50,000 |
| $SNF5^{71-120}$ H2A | 94 | 20 | 125,000,000 | 5,000,000 | 50,000 |
| $SNF5^{195-233}$ WT | 94 | 700 | 20,000,000 | 15,000,000 | 3,500 |
| $SNF5^{195-233}$ H2A | 84 | 700 | 20,000,000 | 15,000,000 | 3,500 |

**Supplemental Table 6:** Yeast strains used in this study. All strains were derived from LH2145.

| Strain | Genotype |
|---|---|
| LH2145 | WT, Mat a from sporulation of BY4743: *ura3∆0 his3∆0 leu22∆0 met15∆0* |
| LH2090 | *ΔQsnf5::kanMX6* |
| LH2971 | *SNF5-TAP-His3MX6* |
| LH2973 | *ΔQsnf5-TAP-His3MX6* |
| LH2974 | $_{HtoA}$*snf5-HIS3* |
| LH2975 | $_{HtoA}$*snf5-TAP-kanMX6* |
| LH2991 | *ADH2::*$P_{ADH2}$*-mCherry-URA3* |
| LH2992 | *ΔQsnf5-kanMX6 ADH2::*$P_{ADH2}$*-mCherry-URA3* |
| LH2993 | $_{HtoA}$*snf5-HIS3 ADH2::*$P_{ADH2}$*-mCherry-URA3* |
| LH3486 | *met15∆0 SNF5::kanMX6 (CEN/ARS-SNF5::URA3)* |
| LH3513 | *snf5Δ::kanMX6 ADH2::*$P_{ADH2}$*-mCherry-URA3 (CEN/ARS-SNF5::URA3)* |
| LH3632 | *snf5Δ::kanMX6 ADH2::*$P_{ADH2}$*-mCherry-URA3 TRP1::pHluorin-natMX6 (CEN/ARS-SNF5::URA3)* |
| LH3647 | *ADH2::*$P_{ADH2}$*-mCherry-URA3 snf2::SNF2-TAP-His3MX6* |
| LH3649 | *ΔQsnf5-HIS3 ADH2::*$P_{ADH2}$*-mCherry-URA3 snf2::SNF2-TAP-kanMX6* |
| LH3652 | $_{HtoA}$*snf5-HIS3 ADH2::*$P_{ADH2}$*-mCherry-URA3 snf2::SNF2-TAP-kanMX6* |
| LH3705 | *SNF5 ADH2::*$P_{ADH2}$*-mCherry-URA3 leu2::pHluorin-LEU2* |
| LH3707 | *ΔQsnf5::kanMX6 ADH2::*$P_{ADH2}$*-mCherry-URA3 leu2::pHluorin-LEU2* |
| LH3713 | $_{HtoA}$*snf5-HIS3 ADH2::*$P_{ADH2}$*-mCherry-URA3 leu2::pHluorin-LEU2* |

**Supplemental Table 7:** Plasmids used in this study.

| Plasmid | Identity |
|---|---|
| pLH226 | pFA6a- *ΔQsnf5*-GFP(S65T)-KANMX6 |
| pLH416 | pFA6a-*SNF5*-GFP-KANMX6 |
| pLH887 | pRS316-SNF5 (CEN/ARS plasmid) |
| pLH931 | pFA6a-$_{4HtoA}$*snf5*-KANMX6 |
| pLH963 | pFA6a-*SNF5*-TAP-KANMX6 |
| pLH964 | pFA6a-*SNF5*-TAP-HIS3MX6 |
| pLH998 | pRS306-$P_{ADH2}$-*mCherry* |
| pLH1085 | pFA6a-$_{6HtoA}$*snf5*-HIS3MX6 |
| pLH1093 | pFA6a-*3'snf2*-TAP-KANMX6 |
| pLH1097 | pRS305-$P_{TDH3}$-pHluorin |
| pLH1206 | pFA6a-*3'snf2*-TAP-NATMX |
